# DAAM1 formin and Ena/VASP proteins assemble functionally distinct actin filaments for focal adhesions

**DOI:** 10.64898/2026.08.19.745742

**Authors:** Xiang Le Chua, Parijat Biswas, Hugo Wioland, Pekka Lappalainen

## Abstract

Eukaryotic cells contain multiple biochemically distinct actin filament networks, which enable versatile functions of actin in a range of cellular processes. Yet, the mechanisms by which specific actin filament networks are assembled in a common cytoplasm remain elusive. Here, we investigated how functionally distinct actin nanoscale layers, specified by α-actinin and tropomyosin isoforms, Tpm1.6 and Tpm3.2, are assembled at focal adhesions. By combining genetic perturbations with mitochondrial-targeting of actin polymerases, we discovered that DAAM1 formin assembles Tpm3.2-actin filaments, whereas Ena/VASP family proteins polymerize α-actinin cross-linked actin filament bundles at focal adhesions. Consequently, loss of DAAM1 dampened Tpm3.2 protein levels and impaired focal adhesion disassembly, thus phenocopying Tpm3.2-deficient cells. In contrast, Ena/VASP depletion led to defective focal adhesion maturation and loss of α-actinin from focal adhesions. More broadly, our study highlights specific roles for formin and Ena/VASP family proteins in assembling biochemically and functionally distinct linear actin filament arrays in cells.

## INTRODUCTION

The actin cytoskeleton is a complex, dynamic structure that plays a central role in multiple cellular processes, such as migration, morphogenesis, endocytosis, phagocytosis, mechanosensing, and cytokinesis. This remarkable versatility of the actin cytoskeleton stems from the ability of actin monomers to assemble into biochemically, mechanically, and functionally distinct filaments in a common cytoplasm^1^. For example, interphase yeast cells harbor a branched network of actin filaments at the sites of endocytosis and long linear actin filament cables that mediate transport of cargo to growing regions of the yeast cell^2–4^. The branched actin filaments facilitating endocytic internalization are nucleated by the Arp2/3 complex, which binds to the side of a pre-existing ‘mother’ filament to nucleate a ‘daughter’ filament that elongates as a branch from the mother filament^5^. On the other hand, actin cables are generated by formin-family proteins, which are donut-shaped dimers that associate with actin monomers and the barbed ends of actin filaments to nucleate and polymerize linear, unbranched actin filaments^6–9^. Both structures are composed of specific sets of other actin-binding proteins. For example, in cables, the filaments are covered by tropomyosins, which facilitate myosin V-mediated transport along the actin filaments, while endocytic actin patches are devoid of tropomyosins but contain other actin-binding proteins such as the actin-severing protein cofilin and actin filament cross-linker fimbrin^1,10–12^. Importantly, actin nucleating machineries that assemble distinct structures in cells compete for shared resources^13–17^.

Animal cells possess even more complex actin cytoskeletons than unicellular yeasts. Although mammalian non-muscle cells typically express only two highly homologous actin isoforms, β- and γ-actin, their actin cytoskeletons are composed of a multitude of different sub-structures that display specific protein compositions and cellular functions^18,19^. Accordingly, animal cells also express a much larger repertoire of actin-associated proteins compared to unicellular yeasts. For example, while yeast cells express only one or two tropomyosins, more than 30 tropomyosin isoforms with specific biochemical and cellular functions can be generated from four tropomyosin genes through alternative splicing in human cells^20,21^. Moreover, whereas yeasts possess only two formins, human cells express 15 different formins^22^, as well as several other proteins that nucleate or polymerize linear actin filaments. These include the three Enabled/vasodilator-stimulated phosphoprotein (Ena/VASP)-family proteins, named VASP, Mena and Evl, which associate with actin filament barbed ends to polymerize linear, unbranched actin filaments *in vitro*^23^. However, why animal cells need such a large repertoire of proteins that produce linear actin filaments, and the possible links between different actin filament polymerizing factors and specific cellular actin filament ‘populations’ have remained largely elusive. For example, while both formins and Ena/VASP proteins can generate seemingly similar linear actin filaments *in vitro*, our understanding of the differences between Ena/VASP- and formin-polymerized actin filaments in cells remains incomplete.

Adding to this complexity, a recent study showed that even individual cellular structures, such as focal adhesions, are composed of several functionally distinct actin filament populations. Interferometric photo-activated localization microscopy (iPALM) revealed that cell-matrix adhesions are composed of at least three vertically organized actin filament nanoscale layers: α- actinin cross-linked filaments forming the uppermost layer, followed by tropomyosin-1.6 (Tpm1.6)-actin filaments slightly below, and tropomyosin-3.2 (Tpm3.2) layer toward the bottom of the focal adhesion^24^. Whereas α-actinin cross-linked actin filaments and Tpm1.6 actin filaments appear to connect the focal adhesion to dorsal stress fibers, and the Tpm1.6-actin filaments are critical for focal adhesion maturation^24,25^, Tpm3.2-actin filaments are important for microtubule-dependent focal adhesion disassembly^24^. However, the mechanism by which these functionally specific actin filament layers are assembled at focal adhesions remains unknown.

Here, we show that two different classes of actin filament assembly factors, formins and Ena/VASP-family proteins, are responsible for the assembly of different actin filament populations for focal adhesions. Loss-of-function and mitochondrial-targeting experiments revealed that DAAM1 and DAAM2 formins polymerize Tpm3.2-decorated actin filaments for focal adhesions, whereas α-actinin cross-linked actin filaments are assembled at focal adhesions by Ena/VASP-family proteins. Thus, our findings identify specific roles for two actin assembly machineries in generating biochemically and functionally distinct linear actin filament arrays in cells.

## RESULTS

### Depletion of DAAM1 and DAAM2 formins phenocopies the loss of tropomyosin-3

Tropomyosins and α-actinin form specific nanoscale layers in focal adhesions^24^ and display mutually exclusive localization patterns along actin filaments *in vitro* (**Figure S1A**). To understand how α-actinin cross-linked actin filaments and tropomyosin-actin filaments are assembled at focal adhesions, we first focused on the role of formins by using U2OS osteosarcoma cells as a model system. Among the formin family proteins, DAAM1 has been implicated in regulating focal adhesion dynamics^26,27^ and was shown to localize to the cortex of mammalian cells^28^ and *Drosophila* egg chamber follicular epithelial cells^29^. By transiently expressing EGFP-DAAM1 in U2OS cells, we confirmed that DAAM1 indeed localizes predominantly to the cell cortex. Interestingly, in cells expressing low levels of EGFP-DAAM1, we also observed clear localization of DAAM1 along cortical stress fibers, and occasional enrichment at focal adhesions (**Figure S1B**).

To determine the potential role of DAAM1 in regulating focal adhesion dynamics and specific actin filament populations, we generated U2OS cell clones deficient in DAAM formins via CRISPR-Cas9. We first generated DAAM1 knockouts (clones #1 and #2), and subsequently depleted DAAM2, its closest homologue, from one of the parental DAAM1 knockout clones (DAAM1/DAAM2 double knockout clones #1A and #1B were generated from the parental DAAM1 knockout clone #1). The successful knockouts were validated by Western blot and next-generation sequencing analyses (**Figure S2A, B**). For all assays, the knockout cells were used within ten passages of sorting to reduce the possibility of phenotypic drift.

Compared to the wild-type U2OS cells, the DAAM1 depleted cells were more elongated and often contained abnormally long tails. These morphological defects appeared slightly more pronounced in the DAAM1/DAAM2 double knockout cells (**Figure 1A, B**). The loss of either DAAM1 or both DAAM formins also led to ∼50 % increase in focal adhesion density (**Figure 1C, D**). Moreover, the focal adhesions in the DAAM knockout cells were located closer towards the cell center as compared to the wild-type cells (**Figure 1C, E**). These defects were consistently observed in different DAAM1 and DAAM1/DAAM2 knockout clones analyzed, indicating that they are not a consequence of off-target effects (**Figure 1**). Interestingly, the abnormalities in cell morphology and focal adhesions in the DAAM knockout cells were similar to the ones previously reported for tropomyosin-3 (Tpm3) knockout cells, where two highly homologous *Tpm3* splice variants, Tpm3.1 and Tpm3.2, were depleted^24^. This observation prompted us to analyze the protein levels of each tropomyosin isoform in the DAAM knockout cells. Western blot analysis of DAAM1 and DAAM1/DAAM2 knockout cells revealed a severe (∼80 %) reduction in the Tpm3.1/Tpm3.2 protein levels, whereas the protein levels of other tropomyosins were either unchanged or only moderately decreased (**Figure 1F**). Quantitative real-time PCR revealed that the mRNA levels of Tpm3.1/3.2 remained unchanged in the DAAM1 knockout and DAAM1/DAAM2 double knockout cells (**Figure 1G**), suggesting that the loss of Tpm3.1/Tpm3.2 is not due to alteration at the transcriptional level. Because recent studies demonstrated that tropomyosins are unstable in the absence of an actin filament template in cells, and targeted to the proteosome^30,31^, it is likely that the loss of DAAM formins results in degradation of Tpm3.1/Tpm3.2 proteins in cells, thereby explaining the similar phenotypes observed in DAAM- and Tpm3-knockout cells. Together, these data show that DAAM family formins are essential in regulating cell morphology and focal adhesion dynamics and reveal a link between DAAM formins and Tpm3-actin filaments.

**Figure 1.**
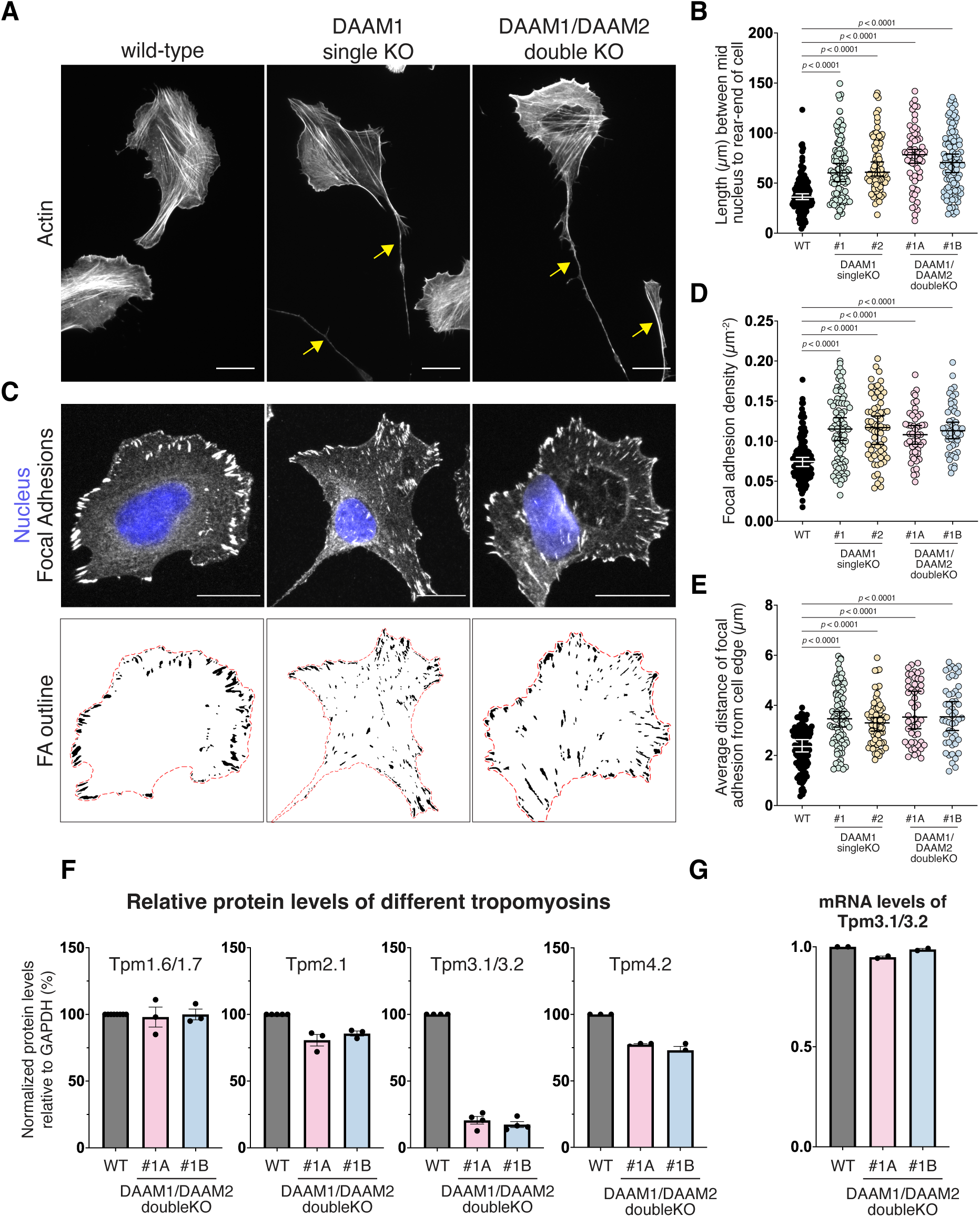
Effects of DAAM1 and DAAM1/DAAM2 depletions on cell morphology, focal adhesions and tropomyosin protein levels. **(A)** Representative micrographs of wild-type, DAAM1 knockout, and DAAM1/DAAM2 double knockout U2OS cells stained for F-actin (phalloidin). The yellow arrows highlight the abnormally long tails of DAAM1 and DAAM1/DAAM2 knockout cells. **(B)** Analysis of tail lengths of wild-type (n = 157 cells), DAAM1 knockout clone #1 (n = 96 cells), DAAM1 knockout clone #2 (n = 79 cells), DAAM1/DAAM2 double knockout clone #1A (n = 60 cells), DAAM1/DAAM2 double knockout clone #1B (n = 102 cells). **(C)** Representative micrographs of wild-type, DAAM1 knockout, and DAAM1/DAAM2 double knockout U2OS cells stained for focal adhesions (vinculin antibody) and nucleus (DAPI). Focal adhesion segmentation and cell outlines (red) are shown below each micrograph. **(D)** Focal adhesion density and **(E)** focal adhesion distributions from the cell edge analyzed in wild-type (n = 127 cells), DAAM1 knockout clone #1 (n = 82 cells), DAAM1 knockout clone #2 (n = 63 cells), DAAM1/DAAM2 double knockout clone #1A (n = 50 cells), DAAM1/DAAM2 double knockout clone #1B (n = 50 cells). **(F)** Relative protein levels of different tropomyosins quantified from cell lysates of wild-type and DAAM1/DAAM2 double knockout clones. Protein levels in wild-type cells were set to 100 %. The data represents mean ± SEM; n ≥ 3 independent cell lysates. **(G)** qRT-PCR for detecting the relative transcript levels of Tpm3.1/3.2 in wild-type and DAAM1/DAAM2 double knockout cells. Error bars in all panels represent mean ± SEM.

### Silencing of Ena/VASP family proteins results in diminished accumulation of α-actinin to focal adhesions and in the loss of focal adhesion -associated dorsal stress fibers

Similar to formins, the Ena/VASP family proteins catalyze the elongation of linear actin filaments at their barbed ends. Ena/VASP proteins also regulate focal adhesions and adhesion-associated dorsal stress fibers^32–35^. Immunofluorescence microscopy with specific antibodies confirmed that in U2OS cells VASP and Mena are enriched at focal adhesions and dorsal stress fibers, where they co-localize with α-actinin (**Figure S3A-D**). Because none of the tested Evl antibodies worked in immunostainings, we studied its localization by expressing mScarlet3-Evl in wild-type U2OS cells. These experiments revealed that Evl also localizes to vinculin-positive focal adhesions in U2OS cells (**Figure S3E**). To elucidate the role of Ena/VASP family proteins in polymerizing actin filaments at focal adhesions, we depleted all members of the Ena/VASP-family from U2OS cells by siRNA. Efficient depletion of VASP (>95 % reduction) and Mena (∼90 % reduction) was confirmed by Western blot (**Figure S4A-C**), whereas the extent of Evl depletion could not be examined due to the lack of specific antibody. Silencing of Ena/VASP proteins resulted in nearly complete loss of dorsal stress fibers, as quantified from cells grown on crossbow-shaped fibronectin micropatterns. In contrast, DAAM1/DAAM2 double knockout cells still exhibited relatively normal quantities of dorsal stress fibers (**Figure 2A-D).** Because dorsal stress fibers elongate through actin polymerization at focal adhesions^36^, we next examined the effects of Ena/VASP silencing on focal adhesions. This revealed that focal adhesions were smaller and located closer to cell edges in the Ena/VASP-silenced cells as compared to control cells (**Figure S5A-D**), providing evidence that Ena/VASP-family proteins are important for focal adhesion maturation.

**Figure 2.**
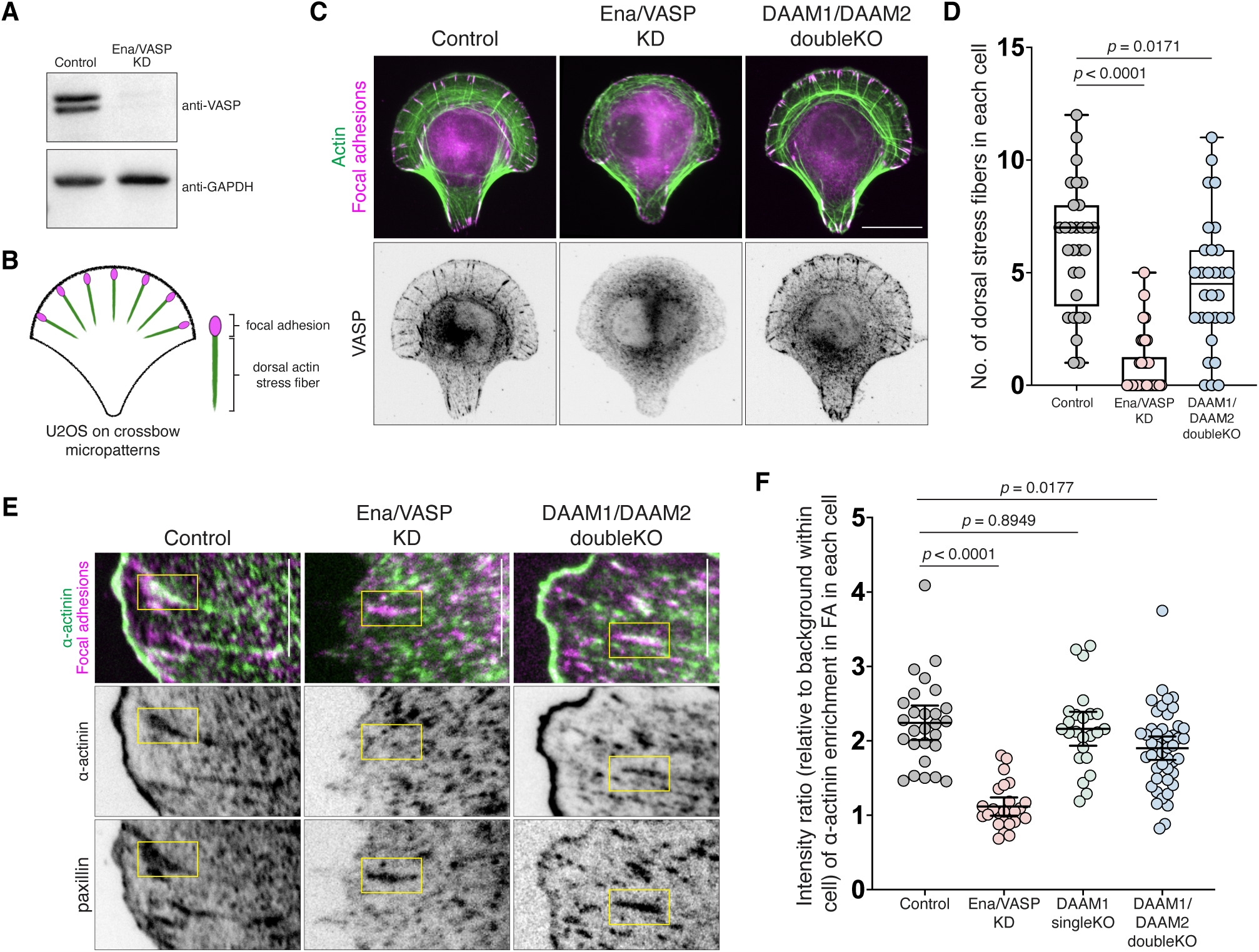
Effects of Ena/VASP depletion on dorsal actin stress fibers and α-actinin localization to focal adhesions. **(A)** Western blot analysis of VASP protein levels in lysates of control cells and in cells treated with siRNAs against Evl, Mena and VASP (Ena/VASP). **(B)** Schematic representation depicting dorsal actin stress fibers and focal adhesions of a wild-type U2OS cell on a fibronectin-coated crossbow micropattern. **(C)** Representative micrographs of control cells, Ena/VASP-depleted cells and DAAM1/DAAM2 double knockout cells on fibronectin-coated crossbow micropatterns stained with anti-vinculin antibody (focal adhesions, FA) and F-actin (phalloidin). Localization of VASP (with specific antibody) is shown below. **(D)** Quantification of the numbers of dorsal stress fibers in control cells (n = 30 cells), Ena/VASP-depleted cells (n = 34 cells) and DAAM1/DAAM2 double knockout cells (n = 30 cells). **(E)** Representative micrographs of control cells, Ena/VASP-depleted cells and DAAM1/DAAM2 double knockout cells seeded on fibronectin-coated conventional coverslips stained with paxillin antibody (focal adhesions, FA) and α-actinin antibody. **(F)** Quantification of the enrichment of α-actinin in paxillin-positive focal adhesions in control cells, Ena/VASP-depleted cells, DAAM1 knockout cells and DAAM1/DAAM2 double knockout cells. Each data point represents the mean value from one cell

Because α-actinin is highly enriched in focal adhesions and dorsal stress fibers^36,37^, we next analyzed the levels of endogenous α-actinin in focal adhesions of Ena/VASP depleted cells and DAAM knockout cells. Interestingly, Ena/VASP silenced cells exhibited a significant decrease in the fluorescence intensity of α-actinin in paxillin-positive focal adhesions, whereas α-actinin accumulation to focal adhesions was not similarly reduced in the DAAM1 knockout and DAAM1/DAAM2 double knockout cells (**Figure 2E, F**). Consistent with earlier studies^35^, the Ena/VASP-silenced U2OS cells migrated with lower speed and exhibited defects in directionality during migration as compared to wild-type cells (**Figure S5E-G**). Considering that, in addition to α-actinin, Tpm1.6 also localizes to dorsal stress fibers^25^, we wondered if the perturbed assembly of dorsal stress fibers in the Ena/VASP-silenced cells would also affect Tpm1.6 or other tropomyosin isoforms. Due to the lack of specific antibodies for detecting specific tropomyosin isoforms by immunofluorescence microscopy, we performed a Western blot analysis on Ena/VASP silenced cells to investigate the protein levels of different tropomyosin isoforms. Interestingly, Ena/VASP-silencing resulted in a severe (∼80 %) reduction in the protein levels of Tpm1.6/Tpm1.7. In contrast, the protein levels of other tropomyosins, including Tpm3.1/Tpm3.2, which was diminished in the DAAM knockout cells, were only moderately affected in the Ena/VASP-silenced cells (**Figure S4D, E**). This result suggests that either the loss of Ena/VASP- polymerized actin filaments, or the resulting depletion of dorsal stress fibers, leads to the degradation of Tpm1.6 in the knockdown cells. Surprisingly, our Western blot analysis also provided evidence that the protein levels of α-actinin were elevated in both Ena/VASP- and DAAM-depleted cells (**Figure S4F, G**). This result seems paradoxical, but may be due to activation of the MRTF/SRF signaling following perturbation of the actin cytoskeleton^38^. Because α-actinin is a target gene of the MRTF/SRF pathway^39^, its expression may therefore be influenced by the actin cytoskeletal organization.^40,41^

Together, these results highlight the importance of Ena/VASP family proteins in focal adhesion maturation and dorsal stress fiber assembly. Moreover, combined with the DAAM knockout results, these data show that, while DAAM formins are critical for generating Tpm3.2-actin filaments in cells, Ena/VASP family proteins control either directly or indirectly α-actinin- and Tpm1.6-actin filaments in cells.

### Mitochondrially-targeted DAAM1 polymerizes tropomyosin-actin filaments, whereas VASP assembles α-actinin cross-linked actin filaments

To further elucidate the link between DAAM1 formin in the assembly of tropomyosin-actin filaments, and to reveal if the Ena/VASP proteins polymerize either α-actinin- or Tpm1.6-actin filaments, or both, we artificially re-localized DAAM1 and VASP to a cellular structure outside of their reported activity regions. To this end, we generated ectopic expression constructs consisting of the TOMM20 mitochondrial-targeting peptide conjugated to the N-terminus of DAAM1 or VASP. These would artificially anchor the fused protein-of-interest on the outer mitochondrial membrane. Because full-length formins are auto-inhibited, we fused the constitutively active C-terminal fragment of DAAM1, consisting of FH1-FH2-DAD domains, to the TOMM20 mitochondrial-targeting peptide (**Figure 3A**). In the case of VASP, we fused the full-length VASP to the TOMM20 mitochondrial targeting peptide. The fluorescence signal of the transiently expressed TOMM20-DAAM1-mEGFP (hereafter Mito-DAAM1), and mitochondrial-targeted VASP construct resembled mitochondria, validated by a specific mitochondrial marker, ATP5A, in U2OS cells (**Figure S6A, B**). We next evaluated the effectiveness of the mitochondrial-targeted DAAM1 and VASP in polymerizing actin filaments. Phalloidin staining revealed strong mitochondrial accumulation of actin filaments co-localizing with mito-DAAM1 in all cells expressing mitochondrially-targeted Mito-DAAM1, demonstrating that it was sufficient to stimulate actin polymerization on mitochondria (**Figure 3B, C, Figure S6C, D**). However, artificial re-localization of VASP to the mitochondrial outer membrane did not appear to stimulate actin polymerization, as judged by the absence of F-actin signal on mitochondria (**Figure S7A, B**). Addition of a flexible linker sequence GGSGGS between TOMM20 mitochondrial-targeting peptide and full-length VASP did not enhance actin polymerization at mitochondria, indicating that the full-length VASP was inactive on the mitochondrial outer membrane (**Figure S7C**).

**Figure 3.**
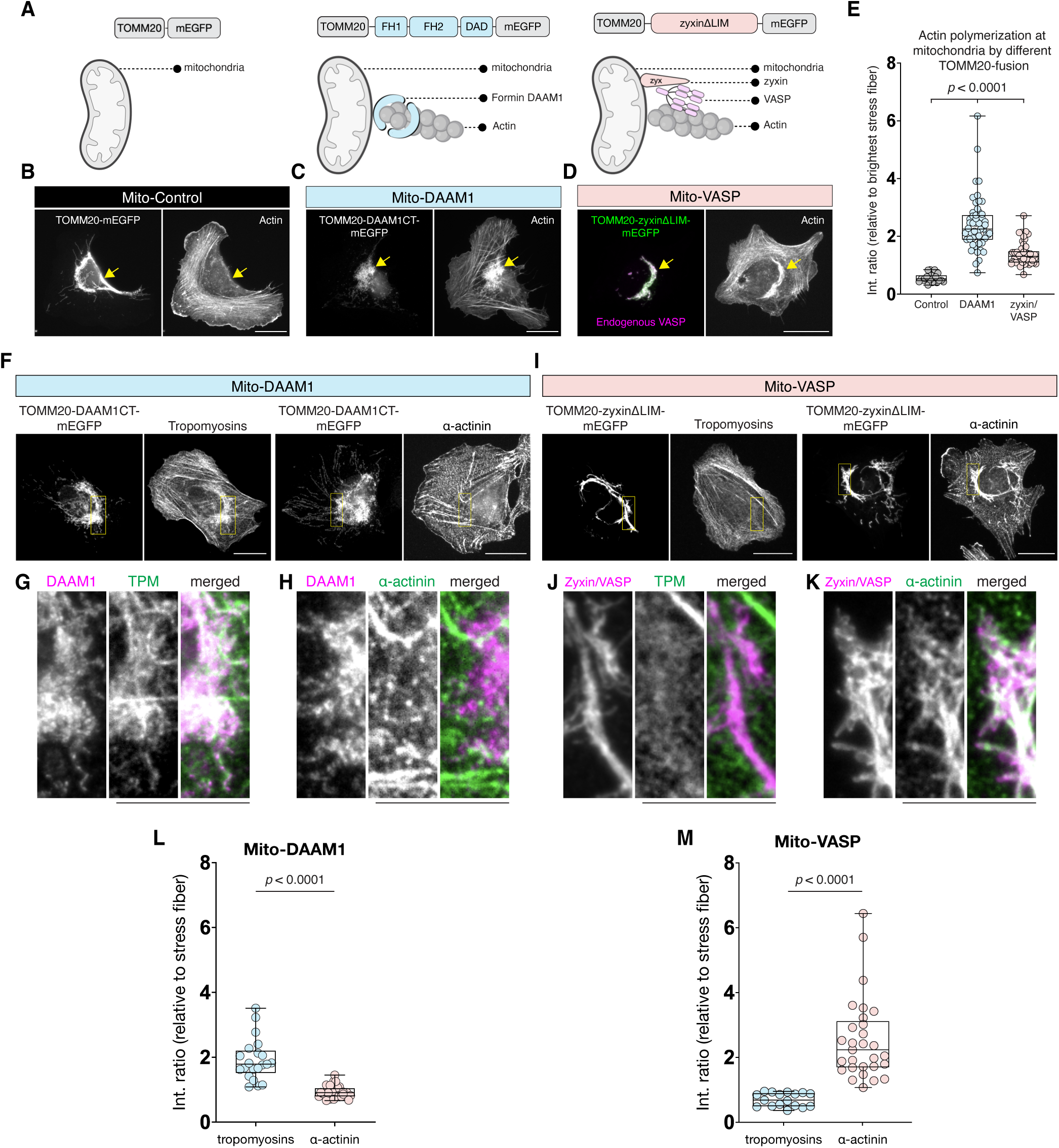
Assembly of tropomyosin-actin and α-actinin-actin filaments following ectopic expression of mitochondrially-targeted DAAM1 and VASP. **(A)** Schematic representation of TOMM20 mitochondrial-targeting peptide fusion-constructs used in the study. **(B-D)** Representative micrographs of wild-type U2OS cells expressing TOMM20-mEGFP (mito-control, panel B), TOMM20-DAAM1CT-mEGFP (mito-DAAM1, panel C) and TOMM20-zyxinΔLIM-mEGFP (mito-VASP, panel D), stained with phalloidin for F-actin and VASP (panel D). **(E)** Quantification of the enrichment of F-actin at the mitochondria in cells expressing mito-control (n = 23 cells), mito-DAAM1 (n = 55 cells) and mito-VASP (n = 46 cells). **(F-H)** Representative micrographs and respective insets of wild-type U2OS cells expressing mito-DAAM1 stained with antibodies against tropomyosin or α-actinin. Yellow boxes in the whole-cell images indicate the regions shown in the insets in panels G and H. **(I-K)** Representative micrographs of wild-type U2OS cells expressing mito-VASP stained with antibodies against tropomyosins or α-actinin. Yellow boxes depict the insets shown in panels J and K. **(L-M)** Quantification of the enrichment of tropomyosin and α-actinin signals at mitochondria of cells expressing mito-DAAM1 (panel L) (n = 21 cells stained for tropomyosin; n = 31 cells stained for α-actinin), and mito-VASP (panel M) (n = 17 cells stained for tropomyosin; n = 29 cells stained for α-actinin). Error bars in all panels represent mean ± SEM. Scale bars for micrographs, 20 µm. Scale bars for insets, 10 µm.

In cells, zyxin, a scaffolding protein, recruits VASP to focal adhesions and is important for the normal morphology and protein composition of focal adhesions^42^. To mimic the context at focal adhesions, we fused the TOMM20 mitochondrial-targeting peptide to the N-terminal fragment of zyxin truncated of its LIM domains, considering that the LIM domains of zyxin exhibit affinity for actin^43^. Artificial re-localization of this zyxin construct to the mitochondrial outer membrane led to an efficient recruitment of endogenous VASP, Mena and Evl on the mitochondria (**Figure 3D, Figure S8A-D**). Importantly, when stained with phalloidin, we observed strong mitochondrial accumulation of actin filaments co-localizing with both VASP and zyxin (**Figure 3D, E**).

Because the mitochondrially-targeted DAAM1 and Ena/VASP (via zyxin) were successful in polymerizing actin filaments on mitochondria, we next examined the possible localization of tropomyosins and α-actinin to the mitochondrial DAAM1- and Ena/VASP-polymerized actin filaments. Interestingly, in cells expressing Mito-DAAM1, we observed a robust accumulation of tropomyosins (as detected with a LC24 antibody recognizing Tpm2.1 and Tpm4.2) on the mitochondria, while α-actinin was not enriched on mitochondria (**Figure 3F-H, Figure S6D**). Conversely, we found that cells expressing Mito-VASP (targeting Ena/VASP proteins to mitochondria via zyxin construct) exhibited intense α-actinin signal on the mitochondria (**Movie S1**), while tropomyosins did not accumulate to actin filaments polymerized on mitochondria through Mito-VASP (**Figure 3I-K**). Since the tropomyosin LC24 antibody used in this experiment was reported to recognize only Tpm2.1 and Tpm4.2 isoforms^44^, we also ectopically expressed mRuby-fusions of Tpm1.6, Tpm2.1, Tpm3.2 and Tpm4.2 in the Mito-VASP transfected cells. While α-actinin still robustly localized on mitochondrial actin filament structures in these cells, none of the tropomyosin isoforms, including Tpm1.6, was enriched on mitochondria (**Figure S8E**). Collectively, these results demonstrate that Mito-DAAM1 and Mito-VASP induce the assembly of biochemically distinct actin filament arrays on the surface of mitochondria, with DAAM1 assembling tropomyosin-actin filaments (**Movie S2-S3**) and VASP assembling α-actinin cross-linked actin bundles (**Movie S1**).

### Mitochondrially-targeted DAAM1 preferentially assembles tropomyosin-3.2-actin filaments

Considering that DAAM1 stimulates the assembly of tropomyosin-decorated actin filaments on the surface of mitochondria, and that the depletion of DAAM1 results in specific degradation of the Tpm3.2 isoform in cells, we further examined the possible link between DAAM1 and Tpm3.2- actin filaments. To this end, we co-expressed Mito-DAAM1 with individual mRuby-tropomyosin fusion proteins in cells and analyzed the accumulation of tropomyosin fusion proteins to mitochondria. To avoid over-expression artifacts, only cells expressing low levels of Mito- DAAM1 and mRuby-fusions of the tropomyosin isoforms were included in the analysis. Expression of mito-DAAM1 led to the recruitment of Tpm1.6, Tpm2.1, Tpm3.2 and Tpm4.2 to the outer mitochondrial membrane, as judged by the visual appearance and co-localization of both mito-DAAM1 and the corresponding co-expressed tropomyosin isoform. Interestingly, Tpm3.2 appeared to accumulate on mitochondria more strongly as compared to the other isoforms (**Figure 4A**). Quantitative analysis of the imaging data confirmed a significantly stronger enrichment of Tpm3.2 on mitochondria as compared to Tpm1.6, Tpm2.1 and Tpm4.2 in cells co-expressing mito- DAAM1 (**Figure 4B**). These data, together with the DAAM knockout results, provide evidence that DAAM1 assembles Tpm3.2-decorated actin filaments in cells.

**Figure 4.**
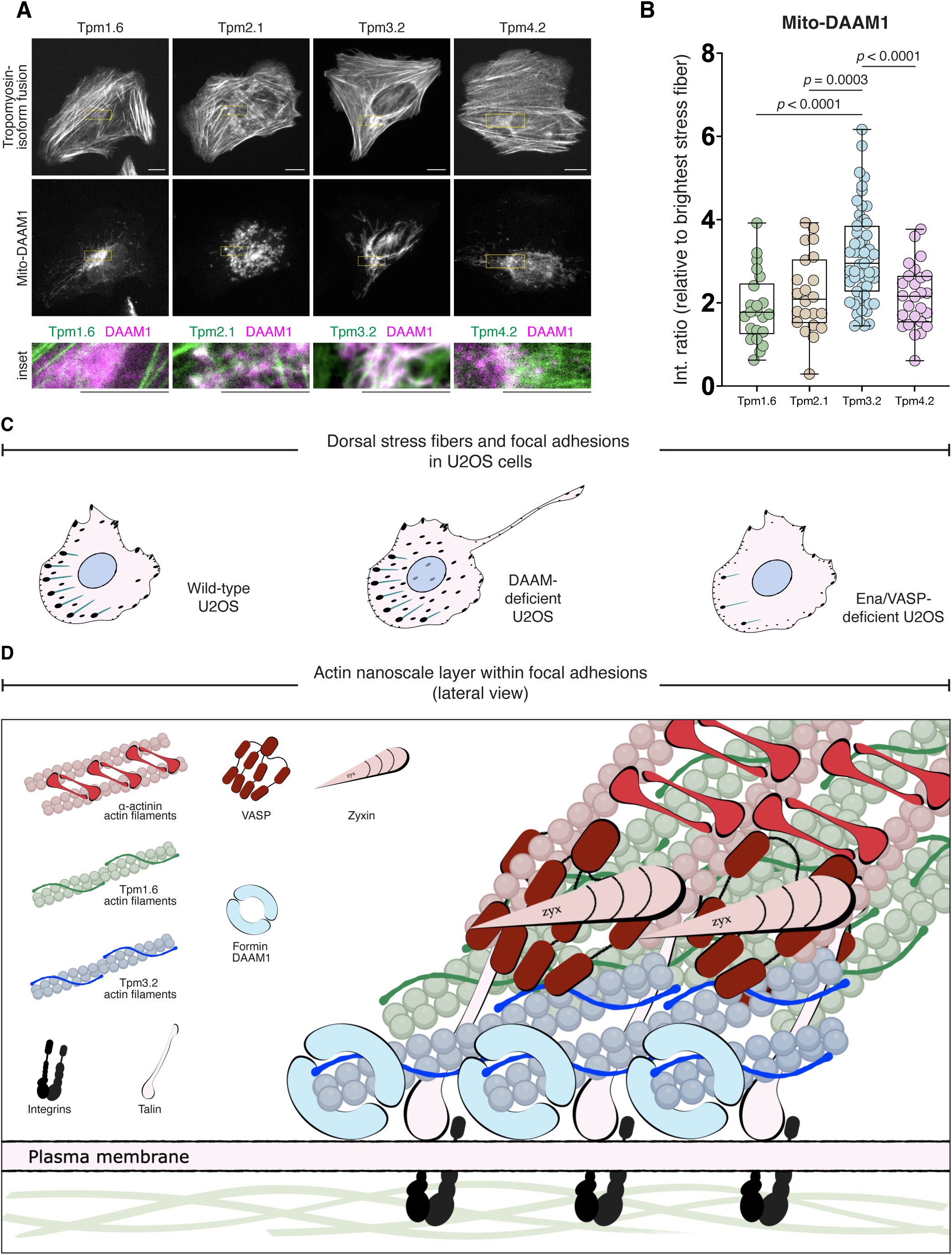
Assembly of specific tropomyosin-actin filaments by mitochondrial-targeted DAAM1. **(A)** Representative micrographs of wild-type U2OS cells co-expressing TOMM20-DAAM1CT-mEGFP (mito-DAAM1) and mRuby-tagged Tpm1.6, Tpm2.1, Tpm3.2 or Tpm4.2. Yellow boxes in micrographs depict the regions shown in the insets below. Scale bars, 10 µm. **(B)** Quantifi- cation of the enrichment of Tpm1.6 (n = 22 cells), Tpm2.1 (n = 22 cells), Tpm3.2 (n = 55 cells), and Tpm4.2 (n = 26 cells) signals at mitochondria of cells expressing mito-DAAM1. Error bars represent mean ± SEM. **(C)** Schematic cartoons representing the effects of DAAM-formin and Ena/VASP family protein depletions on focal adhesions, dorsal stress fibers and cell morphology. **(D)** A working model for actin filament assem- bly at focal adhesions. DAAM1 formin nucleates Tpm3.2-actin filaments, which form the most ventral actin filament layer of focal adhesions that is required for proper adhesion disassembly. VASP, recruited to adhesions by zyxin, polymerizes the uppermost α-actinin cross-linked actin filament layer, which links the adhesion to a dorsal stress fiber. These stress fiber-linked actin filaments are important for force-sensitive matura- tion of focal adhesions.

## DISCUSSION

Focal adhesions contain several vertical layers with specific protein compositions and cellular functions^24,37,45,46^. By combining protein depletion approaches with artificial mitochondrial targeting of actin polymerases, we provide evidence that the uppermost layer of focal adhesions, containing α-actinin cross-linked actin filaments, is generated by Ena/VASP-catalyzed actin polymerization, while the Tpm3.2-actin layer toward the bottom is polymerized by DAAM1 formin **(Figure 4C, D)**. This finding is also consistent with the fact that VASP, and its interaction partner zyxin, localize close to the top layer of focal adhesions.^37^ Our mitochondrial-targeting experiments also revealed that full-length VASP alone is not active, at least on the surface of mitochondria, but that it becomes active upon interacting with zyxin. Sequestering of endogenous Ena/VASP to the mitochondrial surface via ActA in a previous study also revealed the lack of observable actin signal on the mitochondria^32^. Thus, it is possible that zyxin not only targets VASP to focal adhesions^42^, but also activates VASP at adhesions. Depletion of Ena/VASP proteins led to defects in focal adhesion maturation and to a nearly complete loss of dorsal stress fibers. These phenotypes are similar to the ones reported for cells lacking Tpm1.6 and Tpm1.7^24^, which localize to focal adhesions and along dorsal stress fibers, similar to α-actinin. Because mitochondrially-targeted VASP did not induce polymerization of Tpm1.6-actin filaments, despite strong accumulation of α-actinin, we propose that the lack of dorsal stress fibers in Ena/VASP-depleted cells indirectly affects the stability of Tpm1.6/Tpm1.7 to induce their degradation.

Depletion of DAAM-family formins resulted in diminished Tpm3.1/Tpm3.2 protein levels and in very similar phenotypes to those previously reported for the cells lacking Tpm3.1 and Tpm3.2^24^. Moreover, mitochondrial-targeting of DAAM1 provided evidence that it preferentially polymerizes Tpm3.2-actin filaments in cells. Previous work^28,29^ and our results here provided evidence that DAAM1 localizes to the cell cortex, with some enrichment at cortical stress fibers and focal adhesions. Thus, it is possible that DAAM1 polymerizes Tpm3.2-actin filaments directly at focal adhesions. Alternatively, the DAAM1-polymerized cortical actin filaments may become concentrated at focal adhesions similarly to the coalescence of cortical actin filaments during the assembly of cortical stress fibers^47^. The latter option is supported by a recent study suggesting that in *Drosophila* egg chamber follicular epithelial cells, cortically localized DAAM formin supplies actin filaments for stress fiber assembly^29^.

How do VASP and DAAM1 formin polymerize functionally specific actin filaments in cells? Because VASP is a tetramer that additionally forms clusters to promote processive actin filament elongation^48,49^, it is well-suited for assembling actin filaments, which become associated with a cross-linking protein. It was shown that actin filament cross-linkers are sorted *in vitro* on different actin filament structures depending on the spacing between the filaments^50^. Thus, we speculate that at focal adhesions, zyxin may activate VASP and control the spacing between the assembled actin filaments such that they form an optimal template for cross-linking by α-actinin. On the other hand, formins nucleate and elongate individual actin filaments, which are not ideal templates for cross-linking proteins such as α-actinin^8,9,51^. Furthermore, whereas Ena/VASP-family proteins preferentially elongate pre-existing actin filaments, formins can also nucleate actin filaments *de novo* leaving the filament pointed end available for other proteins. Thus, it is likely that in addition to VASP and DAAM1, other cellular factors are required for sorting α-actinin and Tpm3.2 between VASP- and DAAM1-polymerized actin filaments. Good candidates include tropomodulins, which directly associate with tropomyosins at filament pointed ends^52,53^. Importantly, depletion of tropomodulins led to a nearly complete loss of tropomyosin-actin filaments in U2OS cells^31^. However, future work is required to identify the entire protein networks that operate with VASP and DAAM1 to sort α-actinin and Tpm3.2 to specific actin filament layers at focal adhesions. Moreover, the molecular mechanism by which DAAM1 formin polymerizes preferentially Tpm3.2 actin filaments over other tropomyosin isoforms remain to be solved. This may require additional formin-associated proteins. Alternatively, the tropomyosin isoform specificity may arise from different polymerization kinetics of specific formins^54–56^ producing actin filaments of different lengths that may favor filament-binding of specific tropomyosin isoforms.

Collectively, by combining knockout/knockdown approaches with artificial localization of proteins to the mitochondria surface, we provide evidence that Ena/VASP-family proteins and DAAM1/DAAM2 formins polymerize α-actinin cross-linked and Tpm3.2-decorated actin filaments, respectively, in cells. These studies also establish a combination of genetic depletion and artificial mitochondrial-targeting of specific actin polymerases as an excellent approach to examine the composition of specific cellular actin filament networks, because reconstituting such complex protein networks *in vitro* can be challenging.

### Limitations of the study

Our attempts towards reconstituting DAAM1- and zyxin/VASP-mediated actin polymerization *in vitro* were unsuccessful, suggesting that the interplay between DAAM1 - Tpm3.2 and VASP - α- actinin most likely involves additional factors. Thus, detailed understanding of the molecular mechanisms by which DAAM1 and VASP generate biochemically and functionally specific actin filaments needs to be further explored *in vitro*.

## Supporting information

Movie S2: Live-cell imaging of mito-DAAM1 and tropomyosin-3.2

## ACKNOWLEDGEMENTS

We thank Thomas Dos Santos and Natalia Nojszewska for purifying proteins for biochemistry experiments, Mirva Tirkkonen for technical assistance, Katharina Ven for advice with Western blots, Maria Fülöp and Maria Vartiainen for insights into MRTF/SRF signaling, as well as Kah Ying Ng and Gemma González Hernádez for advice with qPCR experiments. We also thank Antoine Jégou, Alphée Michelot, Bruce Goode, David Drubin, Guillaume Romet-Lemonne, Johanna Ivaska, Leonardo Almeida-Souza, Reena Kumari, Sara Wickström, and all members of the Lappalainen group for discussions. We acknowledge the Light Microscopy Unit (LMU) of Institute of Biotechnology, Helsinki, for the use of microscopes. We thank Mikko Liljeström and the Biomedicum Imaging Unit (BIU) for their extensive technical support and use of microscopes.

We are grateful for the reagents provided by Peter Gunning and his research group. We thank the Company of Biologist supporting our *in vitro* biochemistry experiments with a Travelling Fellowship JCSTF25051831 (to X.L.C). This study was supported by grants from Research Council of Finland (346133 and 346131) and Sigrid Juselius Foundation (4708344) to P.L.

## AUTHOR CONTRIBUTIONS

P.L. conceptualized and supervised the project. X.L.C., P.B., and P.L. designed the study. P.B. performed most experiments involving siRNA-mediated silencing of Ena/VASP family proteins and live-cell microscopy. H.W. and X.L.C performed the *in vitro* biochemistry experiments.

X.L.C. performed all other experiments and analyzed the results, with contributions from P.B. X.L.C. and P.L. wrote the manuscript with input from other authors.

## RESOURCE AVAILABILITY

### Lead contact

Further information and requests for resources and reagents should be directed to and will be fulfilled by the lead contact, Pekka Lappalainen

### Material availability

Plasmids and cell lines generated in this study are available from the lead contact upon reasonable request.

### Data and code availability

All data reported and codes used for analysis will be shared by the lead author upon request.

## EXPERIMENTAL MODEL AND SUBJECT DETAILS

### Cell culture and transfection

Human osteosarcoma (U2OS) cells (ATCC HTB-96) were maintained in Dulbecco’s modified Eagle’s medium (DMEM) (ECB7501L, Euro Clone) containing high glucose (4.5 g/L), supplemented with 10 % FBS (10500-064, GIBCO), and Pen-Strep-Glutamine solution (10378016, 100X, GIBCO), at 37 °C, 5 % CO_2_ and 95 % relative humidity. The maximum number of passages for wild-type cells was limited to 30. All freshly thawed cell lines used in this study were passaged at least once before starting the first experiment. All cell lines were grown between 10% - 80% confluence, and the growth medium was replaced every two days. All cell lines were routinely tested for mycoplasma contamination using the Mycoalert Mycoplasma Detection Kit (LT07-418, LONZA), and counterstained with DAPI during immunofluorescence staining throughout the course of this study. Transfections were performed with 200-500 ng of plasmid each into a well of 6-well plate containing ∼50,000 wild-type U2OS cells, using the FuGENE HD (Promega) according to manufacturer’s instructions at 3:1 Fugene/DNA ratio (E2312, Promega). Incubation time for transient transfections was 6 h prior to washout into fresh growth medium, and allowed to rest overnight before each experiment. To visualize the live-cell dynamics of DAAM1 formin in cells, only 200 ng of EGFP-DAAM1 was transfected into a well of 6-well plate containing ∼ 50,000 wild-type U2OS cells, and allowed to rest over 72 h before performing TIRF microscopy.

### Plasmids and molecular cloning

All plasmids generated in this study were subcloned via overlap PCR with KAPA HiFi HotStart ReadyMix (KK2601, Roche Sequencing Store) and ligated through Gibson assembly method (all primers used for cloning are listed in Table S1). Plasmid sequences were verified by whole plasmid sequencing at Eurofins sequencing services (Eurofins Genomics, Germany).

### Protein purification

Skeletal muscle actin (UniProt P68135) was purified from rabbit muscle acetone powder following the protocol as described in previous studies^58,59^. Tpm3.1 C154L C170L C226L A47C was co-expressed in *E. coli* with NatA to ensure N-terminal acetylation. Tpm3.1 was fluorescently labeled with Alexa Fluor 568 Maleimide on the only available cysteine, C47. All native cysteines were replaced by leucines to prevent deleterious labeling. Production, purification and labeling were performed as described in previous study^60^. GST-α-actinin-1 was expressed in BL21 DE3 Star and induced overnight at 16°C. The protein was purified on Glutathione Sepharose 4B beads, cleaved by incubating with Precision Protease overnight at 4°C, eluted and further purified on an HiLoad 16/60 Superdex 200 pg column. Finally, the protein was fluorescently labeled with Alexa Fluor 488 Maleimide.

### *In vitro* TIRF experiment

Experiments were performed in “open chambers”, assembled from two ethanol-cleaned coverslips, separated by parallel strips of double-sided tape. Chambers were passivated with BSA (2.5%, 5 min), rinsed with F-buffer (10 mM Tris-Hcl pH7.4, 100 mM KCl, 1 mM MgCL2, 0.2 mM EGTA, 0.2 mM ATP, 10 mM DTT, 2 mM DABCO). Pre-polymerized actin filaments were mixed with Tpm3.1 and α-actinin in F-buffer supplemented with 0.25% methylcellulose and immediately injected inside the flow chamber. Proteins were visualized on a Nikon Ti inverted microscope with TIRF microscopy (Gataca) as detailed in previous study^60^.

### Generation of knockout cell-lines

All CRISPR knockout cell lines reported in this study were generated as described previously^61^. Guide sequence targeting exon 2 of human DAAM1 gene (ATCACCCAGAAATCACGTAT) and human DAAM2 gene (CAGCCCCATCCCGAACGCAG) were selected based on the CRISPR Design tool with the highest target efficiency scores. Oligonucleotides for cloning the respective guide RNA sequences into pSpCas9 (BB)-2A-GFP vector (48138; a gift from F. Zhang, Addgene, Cambridge, MA) were designed as described previously^62^. Transfected cells were sorted with FACSAria II (BD), using low intensity GFP-positive pass gating, as single cells onto a 96-well plate supplemented with 10 mM HEPES and 20 % FBS containing DMEM media. Two CRISPR clones for DAAM1 single knockout and DAAM1/DAAM2 double knockout were selected for this study based on the absence of detectable DAAM1 and DAAM2 proteins by Western blot analysis. All knockouts were further validated by Sanger Sequencing (Eurofins genomics sequencing service) and next generation sequencing (NGS) (Illumina MiSeq). NGS was performed at the DNA Sequencing and Genomics Laboratory (BIDGEN) laboratory (Institute of Biotechnology, University of Helsinki, Finland).

### siRNA experiments

Dharmacon ON-TARGETplus SMARTpool siRNAs were used for knockdown of ENAH (Cat# L-021932-00-0005), VASP (Cat# L-019763-01-0005), and Evl (Cat# L-020877-00-0005), with Qiagen AllStars Negative Control siRNA (Cat# 1027281) as a control. For knockdown experiments, ∼ 50,000 wild-type U2OS cells were seeded on a 6-well plate. After 18 hours, the cells were transfected with 40 nM siRNA for each gene (total 240 pmol siRNA per well) using jetPRIME transfection reagent (#101000015, Polyplus) following the manufacturer’s protocol. Media was replaced 6 hours post-transfection to minimize reagent-induced toxicity. At 72 hours post-transfection, cells were either seeded on fibronectin-coated glass coverslips for immunofluorescence experiments or lysed for Western blot analysis.

### Western Blotting

All preparations for Western blot analysis were performed on ice. Cell lysates were prepared by washing the cells once with ice-cold PBS and scraping them into lysis buffer (50 mM Tris-HCl pH 7.5 150 mM NaCl, 1 mM EDTA, 10% Glycerol, 1% Triton X-100) supplemented with 1 mM PMSF, 10 mM DTT, 40 mg/ml DNase I and 1 mg/ml of leupeptin, pepstatin, and aprotinin. Protein concentrations were determined with DC Protein Assay (#5000116, Bio-Rad) and equal amounts of total cell lysates were mixed with Laemmli sample buffer, boiled for 5 mins before running on 4–20 % gradient SDS-PAGE gels (#4561096, Bio-Rad). Proteins were transferred to 0.2 μm nitrocellulose membranes (#1704158, Trans-blot turbo transfer pack, Biorad) using Trans-Blot Turbo transfer system (#1704150, Biorad). Membranes were blocked with 5% skim milk in PBS- T (0.05 % Tween-20) for an hour at room temperature with gentle agitation overnight. After blocking, membranes were incubated with primary antibodies diluted in fresh blocking buffer overnight at 4 °C with gentle agitation. However, all tropomyosin antibodies required membranes to be blocked and incubated with 5% BSA in PBS-T instead. After incubation with primary antibodies, membranes were washed three times with PBS-T (0.05 % Tween-20) before incubation with HRP-conjugated secondary antibodies diluted in fresh blocking buffer at room temperature for an hour with gentle agitation. Proteins bands were visualized using Western lightning ECL pro substrate (NEL122001EA, Revvity) in the ChemiDoc XRS+ System (#1708265, Biorad) and quantified using Image Lab™ (#1709690, Biorad) and Fiji ImageJ 1.54f software. GAPDH was used as a loading control for all Western blot experiments. The following primary antibodies and dilutions were used for Western blotting: anti-DAAM1 (1:1000); anti-DAAM2 (1:500); anti-GAPDH (1:7500); anti-TM311 for detecting Tpm1.6/1.7 (1:1000); anti-γ9d for detecting Tpm3.1/3.2 (1:200); anti-LC24 for detecting Tpm2.1 and Tpm4.2 (1:500); anti- δ9d for detecting Tpm4.2 (1:500).

### Real-time quantitative PCR

Total mRNAs were extracted with (Z6014, Promega) and single-stranded cDNA was synthesized (#M0253, NEB) from 500 ng of the extracted mRNA. Primer sequences for amplifying region of Tpm3.2 and GAPDH were taken from previous study^31^. Quantitative PCR reactions were performed with Maxima SYBR Green/ROX (K0221, Thermo Fisher Scientific) in LightCycler 480 Instrument II (Roche). Changes in expression levels were calculated with the 2^−ΔΔCt^ method and normalized to GAPDH (ΔCt) and wild-type expression levels, respectively.

### Micropatterns

CYTOOchips^TM^ crossbow micropattern coverslips (10-600-10-18, Cytoo) used for this experiment were coated with fibronectin (10 µg/mL) at room temperature for an hour. The fibronectin-coated crossbow micropattern coverslips were washed three times with PBS before seeding cells on them. Seeded cells were allowed to attach onto the coverslips for 3 h before 4 % PFA fixation and immunofluorescence staining.

### Immunofluorescence staining

For knockout studies, wild-type and respective knockout cell lines were seeded on 10 µg/mL fibronectin-coated (Sigma-Aldrich, L2020) coverslips or on 35 mm imaging dishes (Ibidi µ-dish high), at 60 % confluence for 9 h before fixation. Cells were fixed with 4% PFA in PBS at room temperature for 12 min and washed gently with pre-warmed PBS for three times before permeabilized with 0.2% Triton X-100 in PBS at RT for 7 min. Cells on coverslips were blocked with 5 % BSA in PBS for 30 min before incubation with primary antibodies diluted in 5 % BSA (1:200 dilution) for either 1 h at room temperature or overnight at 4 °C in a humidified chamber to prevent drying. Coverslips were washed gently with PBS for three times before incubating with secondary antibodies (1:250 dilution) in PBS for 1 h at room temperature in a humidified chamber. Whenever needed, phalloidin (1:700 dilution) and DAPI (1:1000 dilution) are diluted in PBS together with the secondary antibodies. Coverslips were washed gently with PBS for three times and mounted onto microscopes slides with ProLong™ Glass Antifade Mountant (P36980, Invitrogen).

### Widefield microscopy

Immunofluorescence samples were imaged using DM6000B fully motorized upright fluorescence microscope equipped with a 63x/1.40-0.60 HCX PL APO Lbd.bl. Oil wd=0.10 objective and following optical filters: DAPI-5060C (ex 377/50, em 447/60), CFP-2432C (ex 438/24, em 483/32), GFP-4050B (ex 466/40, em 525/50), TRITC-B (ex 543/22, em 593/40), LED-mCherry-A (ex 578/21, em 641/75), Cy5-4040C (ex 628/40, em 692/40) (IDEX Health & Science, LLC, USA). The images were acquired using Hamamatsu Orca-Flash4.0 V2 sCMOS camera with 2048 × 2048 pixels image resolution. All images were acquired with the same acquisition settings of 250 milliseconds exposure throughout the course of study. Acquired images were processed with Fiji-ImageJ for data analysis.

### TIRF live-cell microscopy

Wild-type cells transfected with EGFP-DAAM1 were seeded on fibronectin-coated 35 mm glass bottom dishes (81158, Ibidi) containing 10 % FBS, glutamax and 25 mM HEPES, 2 hours before imaging and maintained at 37°C, 5 % CO_2_, and 95 % relative humidity throughout the experiment. TIRF live-cell microscopy was performed with The Zeiss Elyra7 Lattice SIM at TIRF mode. The ‘green’ and ‘red/far-red’ fluorescence emissions were split into two pre-aligned Hamamatsu C15440-20UP CMOS cameras using a dichroic beam splitter. Timelapse TIRF videos were taken at time intervals of 10-60 s.

### Random cell migration assay

To prevent cells colliding with each another during imaging, ∼5000 cells were seeded on each well of a fibronectin-coated 12-well plate. Cells were allowed to attach for 3 h before imaging with a Zeiss Cell Discoverer 7 microscope using a 20x objective with 0.5x magnification in Phase Gradient Contrast mode. Imaging was performed with 25 mM HEPES (pH 7.4) supplemented cell culture media, in 5% CO_2_ and at 37 °C. Images were acquired at 15 min time interval for a total of 10 hours. Cell trajectories were extracted by an in-house Fiji/ImageJ macro that stores coordinates from manual mouse-clicks on the nuclei centroids at every frame. Mitotic or colliding cells were excluded from the analysis. The cell trajectories were later used for calculating migration speed *D*⁄*T* and directionality (*d*⁄*D*), where D = total distance travelled, T = time elapsed during travel, and d = distance between the start-point and finish.

### Mitochondrial-targeting assay

The TOMM20*-SBP-mEGFP was a gift from Juan Bonifacino^57^ (Addgene plasmid #120173). All plasmids for targeting each protein-of-interest to the mitochondrial outer membrane were generated by Gibson assembly of PCR products by replacing SBP from TOMM20*-SBP-mEGFP with the protein-of-interest. Primers used for cloning are listed in Supplementary Table S1. Cells transfected with mitochondrial-targeting constructs were seeded on fibronectin-coated coverslips and allowed to attach for 4.5 h before fixation with 4 % PFA. All data reported in this assay were acquired from cells expressing low levels of mitochondrial-targeted constructs. Widefield images were acquired at short excitation time of 250 milliseconds. Post-acquisition image analyses were performed Fiji and plotted by Prism 10 (GraphPad Software, Inc, La Jolla, CA). To quantify the enrichment of actin, α-actinin, or different tropomyosin isoforms induced by the mitochondrial-targeted constructs at the outer mitochondrial membrane, a 10 × 10–pixel ROI was placed on the brightest mitochondrial tubule in the respective actin, α-actinin, or tropomyosin channel mean fluorescence intensities were measured. The measured intensity was then normalized to one of the brightest stress fiber within a 10 × 10–pixel ROI.

### Tail length analysis

Phalloidin and DAPI staining were used to visualize cell morphology and nuclei, respectively. Images taken with phalloidin and DAPI staining were merged to generate dual-color images. Tail length was measured by drawing a ‘freehand’ line on each cell of the merged image from the approximate midpoint of the nucleus to the distal tip of the phalloidin-positive cell tail.

### Quantification of focal adhesion morphometry

Vinculin staining was applied to visualize focal adhesions, and for quantification of focal adhesion density, size and distribution using an in-house image analysis pipeline written in ImageJ/Fiji macro format. The macro is available upon request. Briefly, the algorithm uses a series of scale-space filtering with iterative Gaussian Blur and Top Hat filters to smoothen pixel noise and suppress redundant background intensity. An image filter equivalent to Laplacian of Gaussian is applied to enhance the blob-like focal adhesions and then subjected to unbiased, uniform intensity thresholding for binarizing the image. Finally, using ImageJ’s Analyze Particles function, the focal adhesions are extracted as ROIs for further analysis. Cells that were in contact with neighboring cells were discarded from the analysis. The size range of the focal adhesions to be segmented was constrained between 0.1 to 60 µm^2^ for all experiments to eliminate artifactual objects. Parameters were carefully modified and cross-checked with corresponding images to achieve ideal segmentation. Post-segmentation, the focal adhesion ROIs were used to quantify their individual sizes, number density within the cell’s spread area, and their distances from the cell edge.

### Focal adhesion density

The focal adhesion density per cell on fibronectin-coated coverslips was calculated according to the equation described in ref.^24,63^:

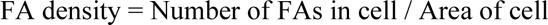

### Quantification of α-actinin in focal adhesions

Sheep anti-paxillin antibody staining was used to visualize focal adhesions instead of vinculin because of host species compatibility with mouse anti-α-actinin and rabbit anti-VASP antibodies which had to be included in this experiment to assess α-actinin enrichment in Ena/VASP depleted cells. To quantify the fluorescence intensity of α-actinin in paxillin-positive focal adhesion in each cell, the measured intensity of α-actinin within each focal adhesion ROI, obtained after focal adhesion segmentation, was normalized to the α-actinin intensity measured within a 2 µm^2^ square ROI positioned in a focal adhesion-free region of the same cell.

### Quantification of dorsal stress fibers density

To quantify the dorsal stress fiber density in cells seeded on fibronectin-coated crossbow micropatterns, the number of dorsal stress fibers was counted along a line parallel to the cell edge and set within of the cell edge. A dorsal stress fiber was defined as an actin bundle with a length of ≥5 µm linked to a focal adhesion at the cell front and pointing towards the cell center.

### Quantification and Statistical Analysis

Statistical tests were performed using two-tailed Students’ t-tests (**Figure 3L, M; Figure S5B-G**), One-way ANOVA followed by Tukey’s multiple comparison’s tests (**Figure 1B-E; Figure 2D, F; Figure 3E**; **Figure 4B; Figure S2G, H; Figure S6E**) using Graphpad Prism. Blind analysis was performed for **Figure 2G**. *P* values < 0.05 was considered statistically significant. Data are presented as mean ± SEM from at least three independent experiments.

**Table S1:**
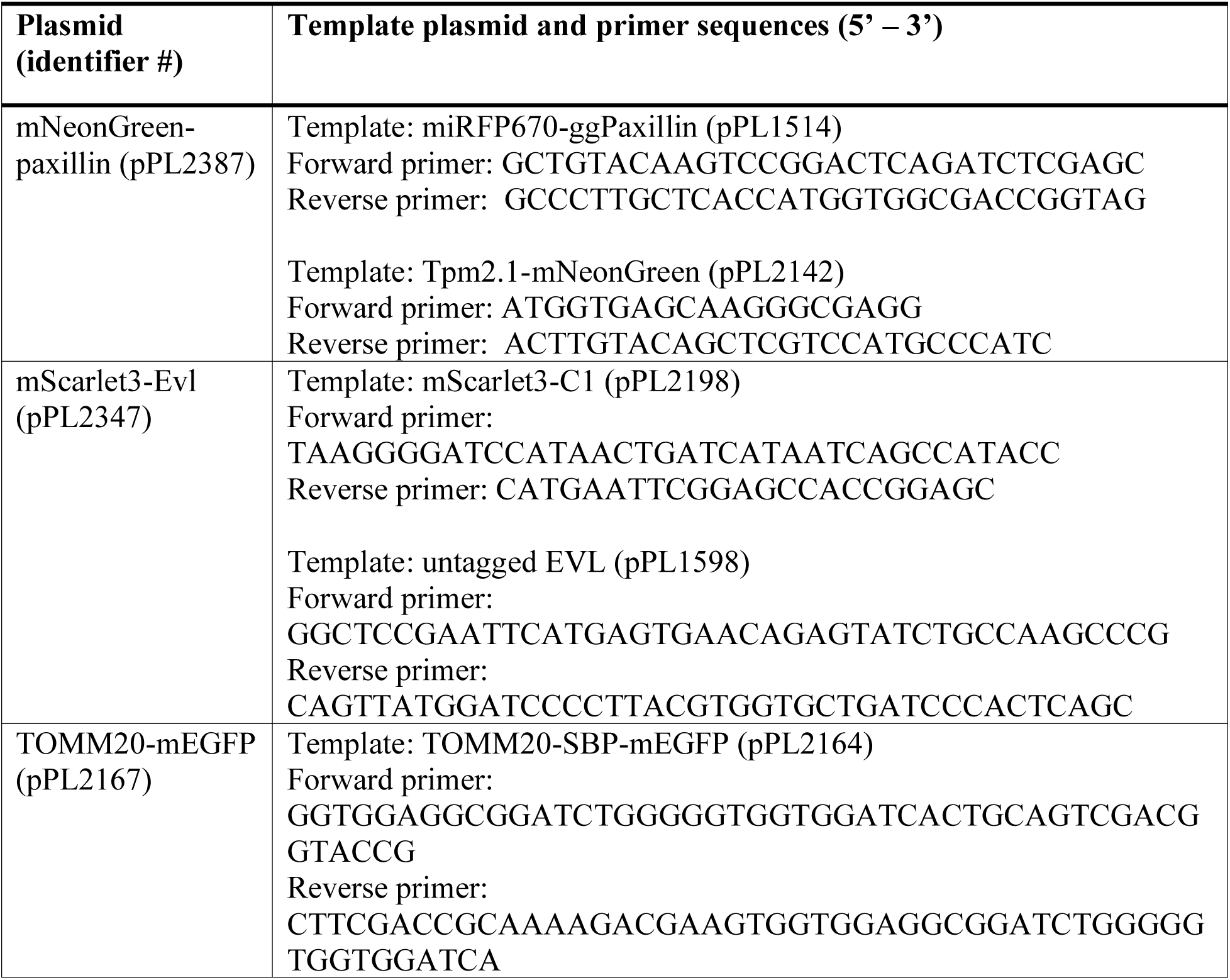
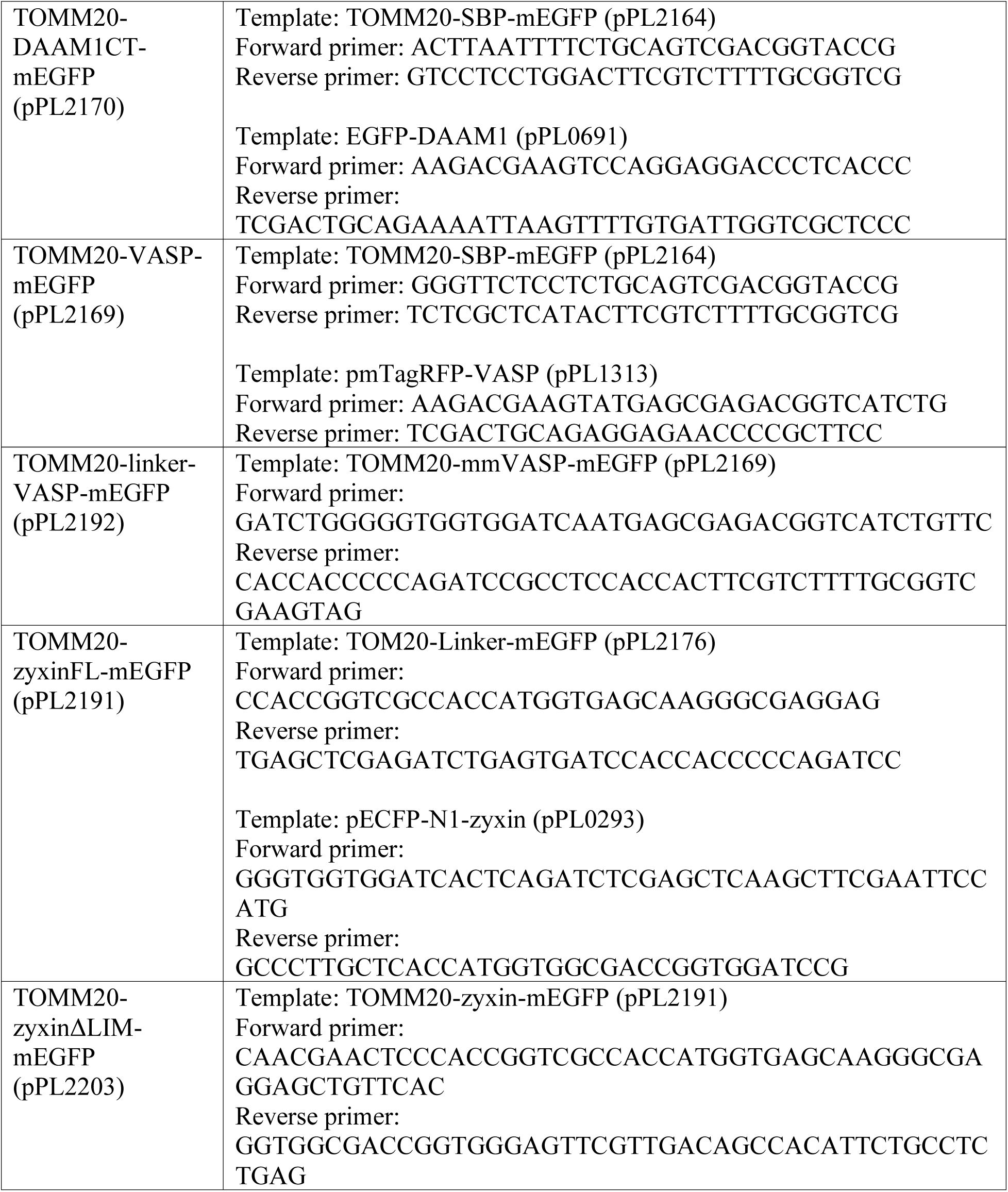
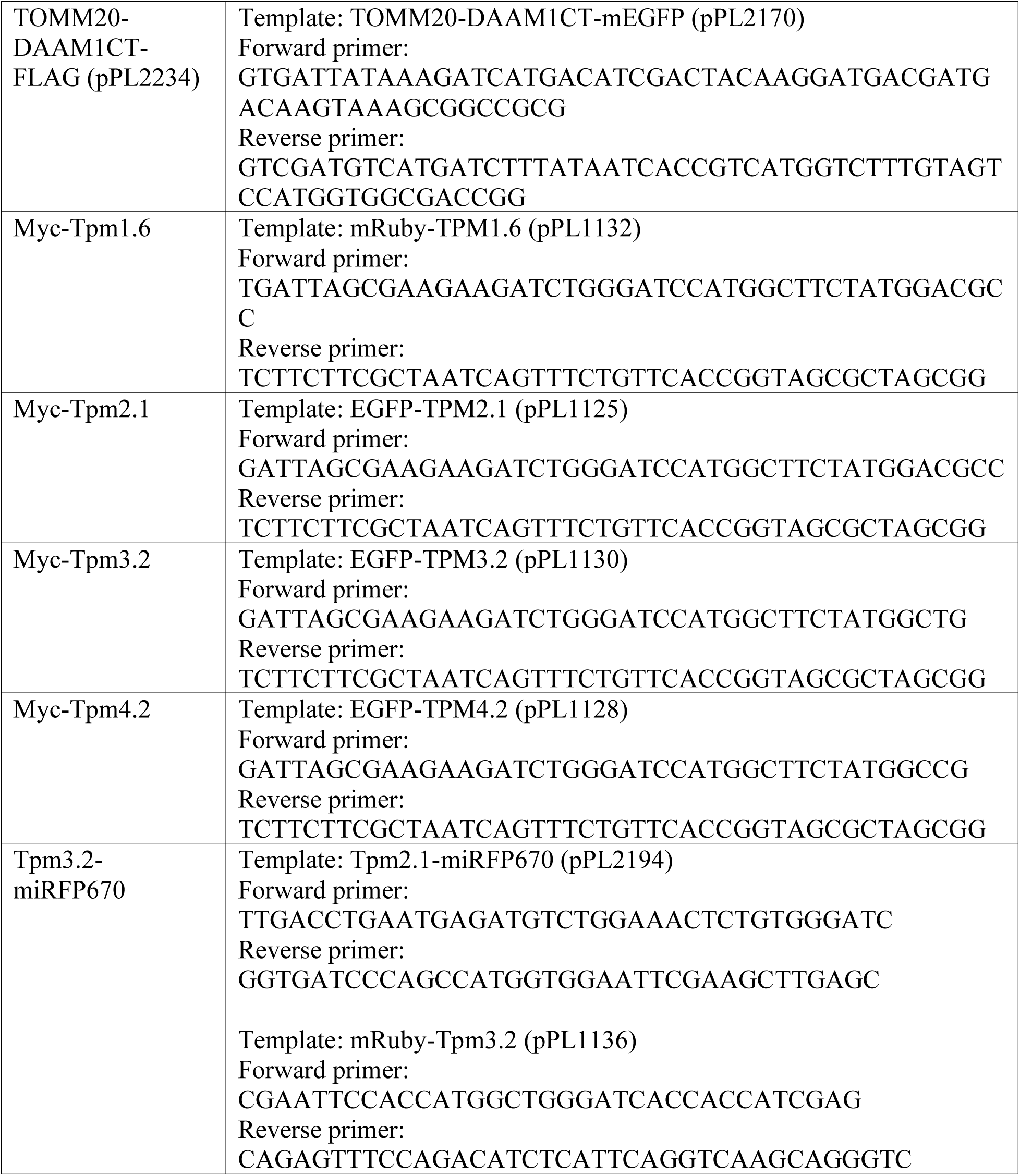
Respective plasmid constructs and primers used in this study.

**Table S2:** Reagents used in this study.

| REAGENT or RESOURCE | SOURCE | IDENTIFIER |
| --- | --- | --- |
| **Antibodies** |  |  |
| Rabbit polyclonal anti-DAAM1 | Proteintech | Cat# 14876-1-AP |
| Rabbit polyclonal anti-DAAM2 | antibodies.com | Cat# A15717 |
| Rabbit polyclonal anti-GAPDH | Sigma-Aldrich | Cat# G9545 |
| Mouse monoclonal anti-vinculin | Sigma-Aldrich | Cat# V9131 |
| Mouse monoclonal anti-TM311 | Sigma-Aldrich | Cat# T2780 |
| Mouse monoclonal anti-γ9d | Peter Gunning | N/A |
| Rabbit polyclonal anti-δ9d | Peter Gunning | N/A |
| Mouse monoclonal anti-LC24 | Peter Gunning | N/A |
| Mouse monoclonal anti-paxillin | BD Bioscience | Cat# 610569 |
| Mouse monoclonal anti-α-actinin-1 | Sigma-Aldrich | Cat# A5044 |
| Rabbit monoclonal anti-VASP | Cell Signaling Technology | Cat# 3132 |
| Sheep polyclonal anti-paxillin | R&D Biosystems | Cat# AF4259 |
| Mouse monoclonal anti-Mena | BD Bioscience | Cat# 610693 |
| Mouse monoclonal anti-ATP5A | Abcam | Cat# 15H4C4 |
| Mouse monoclonal anti-FLAG | Sigma-Aldrich | Cat# F1804 |
| **Bacterial and virus strains** |  |  |
| *E. coli* XL10 gold ultracompetent cells | This study | N/A |
| **Chemicals, peptides, and recombinant proteins** |  |  |
| Alexa Fluor<sup>TM</sup> 488, 568, 647 conjugated to phalloidin | Thermo Fisher Scientific | Cat# A12379, Cat# A1238, and Cat# A22287 |
| Secondary antibodies conjugated to Alexa Fluor<sup>TM</sup> 488, 568 and 647 | Thermo Fisher Scientific | Cat# A11001, Cat# A11004, Cat# A31571, Cat# A11034, and Cat# A21245 |
| 4',6-diamidino-2-phenylindole, dihydrochloride (DAPI) | Thermo Fisher Scientific | Cat# D1306 |
| HRP-conjugated goat anti-rabbit | Thermo Fisher Scientific | Cat# G21234 |
| HRP-conjugated goat anti-mouse | Thermo Fisher Scientific | Cat# 31430 |
| Fibronectin | Merck | Cat# 1080938001 |
| Dulbecco's modified Eagle's medium (DMEM) | Euro Clone | Cat# ECB7501L |
| Fetal Bovine Serum (FBS) | GIBCO | Cat# 10500-064 |
| 10 U/ml penicillin, 10 mg/ml streptomycin, and 20 mM L-glutamine | GIBCO | Cat# 10378-016 |
| **Critical commercial assays** |  |  |
| Mycoalert Mycoplasma Detection Kit | LONZA | Cat# LT07-418 |
| FuGENE HD | Promega | Cat# E2312 |
| jetPRIME transfection reagent | Polyplus | Cat# 101000015 |
| PureLink Genomic DNA Mini Kit | Thermo Fisher Scientific | Cat# K182001 |
| NucleoSpin Gel and PCR Clean-up | Macherey-Nagel, Vingmed | Cat# 740609.250 |
| NucleoSpin Plasmid EasyPure | Macherey-Nagel, Vingmed | Cat# 740727.250 |
| NEBuilder kit | New England BioLabs | Cat# E5520S |
| 4%–20% gradient SDS-PAGE gels | Bio-Rad | Cat# 4561096 |
| Trans-Blot Turbo transfer pack Mini Format | Bio-Rad | Cat# 170-4158 |
| ECL pro substrate | Revvity | Cat# NEL122001EA |
| **Experimental models: Cell lines** |  |  |
| Human osteosarcoma (U2OS) cells | Previous study<sup>24</sup> | N/A |
| **Oligonucleotides** |  |  |
| Primers for cloning, sequencing and sequences of the siRNAs | This study | Table S1 |
| **Recombinant DNA** |  |  |
| ENAH (Mena) SMARTpool siRNA | Dharmacon | Cat# L-021932-00-0005 |
| VASP SMARTpool siRNA | Dharmacon | Cat# L-019763-01-0005 |
| Evl SMARTpool siRNA | Dharmacon | Cat# L-020877-00-0005 |
| AllStars Neg. Control siRNA | Qiagen | Cat# 1027281 |
| pSpCas9(BB)-2A-GFP-DAAM1 | This study | pPL1919 |
| pSpCas9(BB)-2A-GFP-DAAM2 | This study | pPL2020 |
| EGFP-DAAM1 | Gift from Alexander Bershadsky | pPL0691 |
| miRFP670-paxillin | Previous study<sup>24</sup> | pPL1514 |
| mNeonGreen-paxillin | Subcloned from miRFP670-paxillin by replacing miRFP670 with mNeonGreen | pPL2387 |
| mScarlet3-Evl | Subcloned from untagged Evl into mScarlet3-C1 vector | pPL2347 |
| TOMM20-SBP-mEGFP | Gift from Juan Bonifacino (Addgene #120173)<sup>57</sup> | pPL2164 |
| TOMM20-mEGFP (Mito-Control) | Subcloned from TOMM20-SBP-mEGFP by replacing SBP with GGSGGS linker | pPL2176 |
| TOMM20-DAAM1CT-mEGFP (Mito-DAAM1) | Subcloned from TOMM20-SBP-mEGFP by replacing SBP with FH1-FH2-DAD of DAAM1 | pPL2170 |
| TOMM20-VASP-mEGFP | Subcloned from TOMM20-SBP-mEGFP by replacing SBP with VASP | pPL2191 |
| TOMM20-linker-VASP-mEGFP | Subcloned from TOMM20-VASP-mEGFP | pPL2192 |
| TOMM20-zyxinΔLIM-mEGFP (Mito-VASP) | Subcloned from TOMM20-SBP-mEGFP by replacing SBP with zyxin with LIM domains | pPL2203 |
| TOMM20-DAAM1CT-FLAG | Subcloned from TOMM20-DAAM1CT-mEGFP by replacing mEGFP with FLAG | pPL2234 |
| mRuby-Tpm1.6 | Previous study<sup>24</sup> | pPL1132 |
| Myc-Tpm1.6 | Subcloned from mRuby-Tpm1.6 by replacing with mRuby with myc tag. | pPL2173 |
| mRuby-Tpm2.1 | Previous study<sup>21</sup> | pPL1131 |
| Myc-Tpm2.1 | Subcloned from mRuby-Tpm2.1 by replacing with mRuby with myc tag. | pPL2174 |
| mRuby-Tpm3.2 | Previous study<sup>24</sup> | pPL1136 |
| Myc-Tpm3.2 | Subcloned from mRuby-Tpm3.2 by replacing with mRuby with myc tag. | pPL2172 |
| Tpm3.2-miRFP670 | Subcloned from Tpm2.1-miRFP670 (unpublished study) by replacing Tpm2.1 with Tpm3.2 | pPL2195 |
| mRuby-Tpm4.2 | Previous study<sup>21</sup> | pPL1134 |
| Myc-Tpm4.2 | Subcloned from mRuby-Tpm4.2 by replacing with mRuby with myc tag. | pPL2175 |
| TagRFP- α-actinin | Gift from Michael Davidson (Addgene #58033) | pPL1423 |
| **Software and algorithms** |  |  |
| Fiji (Image J) | National Institutes of Health, NIH | https://imagej.net/software/fiji/ |
| Geneious Prime | Biomatters Limited | N/A |
| GraphPad Prism 10 | N/A | https://www.graphpad.com/ |
| Origin | OriginLab Corporation | N/A |

---

**Figure S1.**
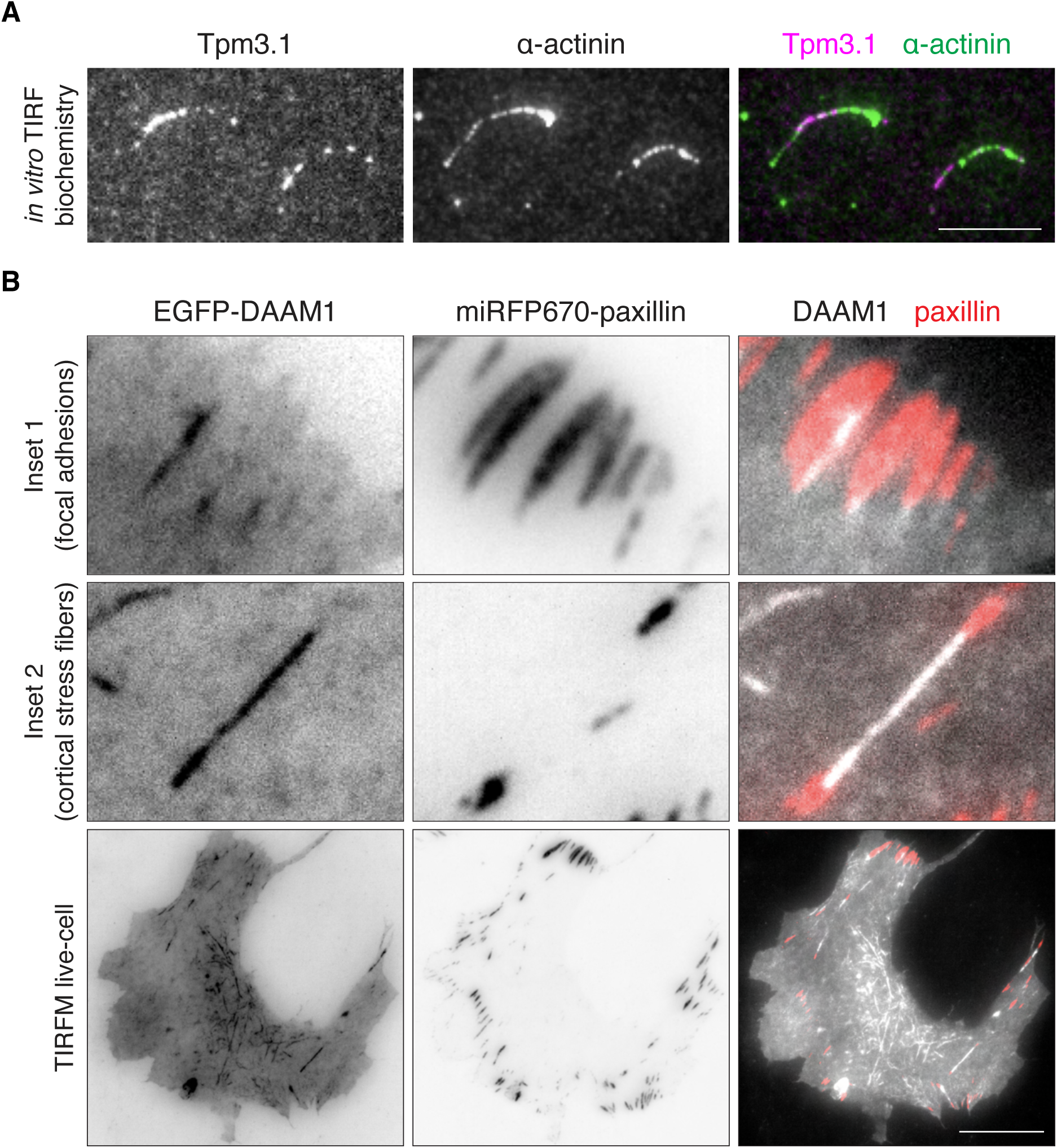
Total internal reflection microscopy (TIRF) demonstrating segregation of purified Tpm3.1 and α-actinin-1 on actin filaments in vitro, and localization of DAAM1 formin in U2OS cells. (**A)** Representative micrographs showing the distributions of Tpm3.1 and α-actinin-1 along 3% ATTO-647 labeled actin filaments (1 mM) in vitro. Tpm3.1 (200 nM, monomeric) and α-actinin-1 (100 nM, monomeric) were labeled with Alexa-561 and -488, respectively. Scale bars, 10 µm. **(B)** Representative micrographs of wild-type U2OS cells co-expressing miRFP670-paxillin and EGFP-DAAM1. Box in red (inset 1) shows an occasional localization of DAAM1 to focal adhesions. Box in blue (inset 2) shows the localization of DAAM1 to cortical stress fibers. Scale bars, 20 µm.

**Figure S2.**
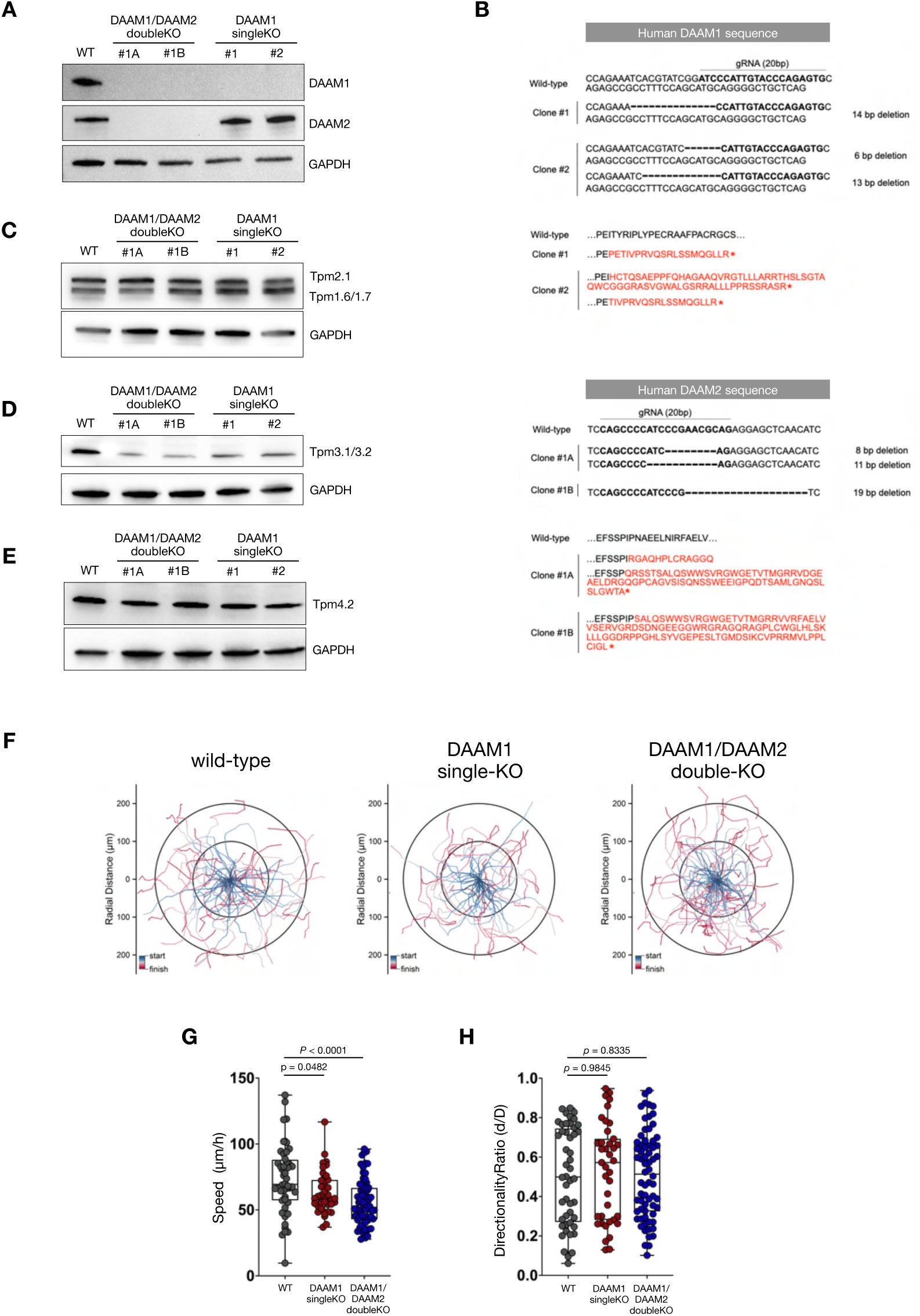
Genomic validation and phenotypes of DAAM knockouts. **(A)** Western blot analysis of DAAM1 and DAAM2 proteins in lysates from parental wild-type, DAAM1/DAAM2 double knockout clones #1A and #1B, and DAAM1 knockout clones #1 and #2. DAAM1 knockout clone #1 was applied to generate double knockout clones #1A and #1B. **(B)** DNA and amino acid sequence alignments surrounding the cutting site in exon 2 of reference human DAAM1 and DAAM2 genes against all knockout clones. Frameshift mutations resulting in premature stop codons (denoted *) are shown in red. **(C-E)** Western blots of protein levels of Tpm1.6/1.7 and Tpm2.1 (as detected by TM311 antibody, panel C), Tpm3.1/3.2 (detected by γ9d antibody, panel D), and Tpm4.2 (detected by δ9d antibody, panel E) in parental wild-type, DAAM1/DAAM2 double knockout clones #1A and #1B, and DAAM1 knockout clones #1 and #2. **(F)** Random migration trajectories of wild-type cells, DAAM1 knockout clones #1 and #2, as well as DAAM1/DAAM2 double knockout clones #1A and #1B showing polar coordinates with the start point synchronized at origin (0,0) and color-coded to indicate the track’s start (blue) and finish (red) after 5 h. **(G-H)** Quantification of cell migration speed (panel G) and directionality ratio (panel H) of the cells. Each data point represents an individual cell. Error bars represent mean ± SEM.

**Figure S3.**
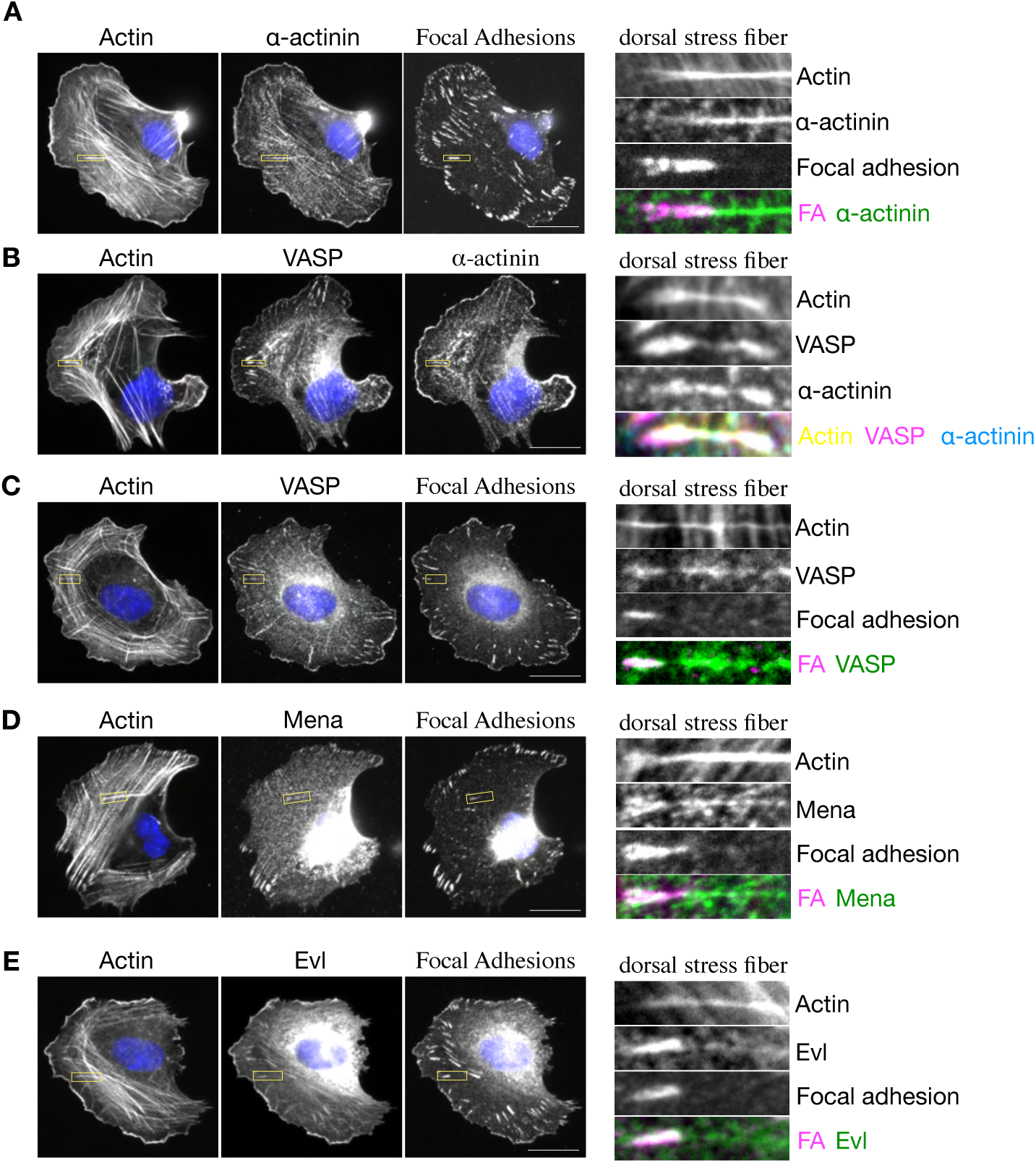
Localization of Ena/VASP family proteins and α-actinin in dorsal stress fibers and focal adhesions. **(A-D)** Representative micrographs of wild-type U2OS cells stained for nucleus (DAPI), F-actin (phalloidin), focal adhesions (paxillin antibody), α-actinin, VASP and Mena (with specific antibodies). Yellow boxes in the whole cell images indicate regions shown in the insets on right. **(E)** Representative micrographs of wild-type U2OS cells expressing mScarlet3-Evl and stained for focal adhesions (vinculin antibody) and F-actin (phalloidin). Yellow boxes in the whole cell images indicate regions shown in the insets on right. Scale bars, 20 µm.

**Figure S4.**
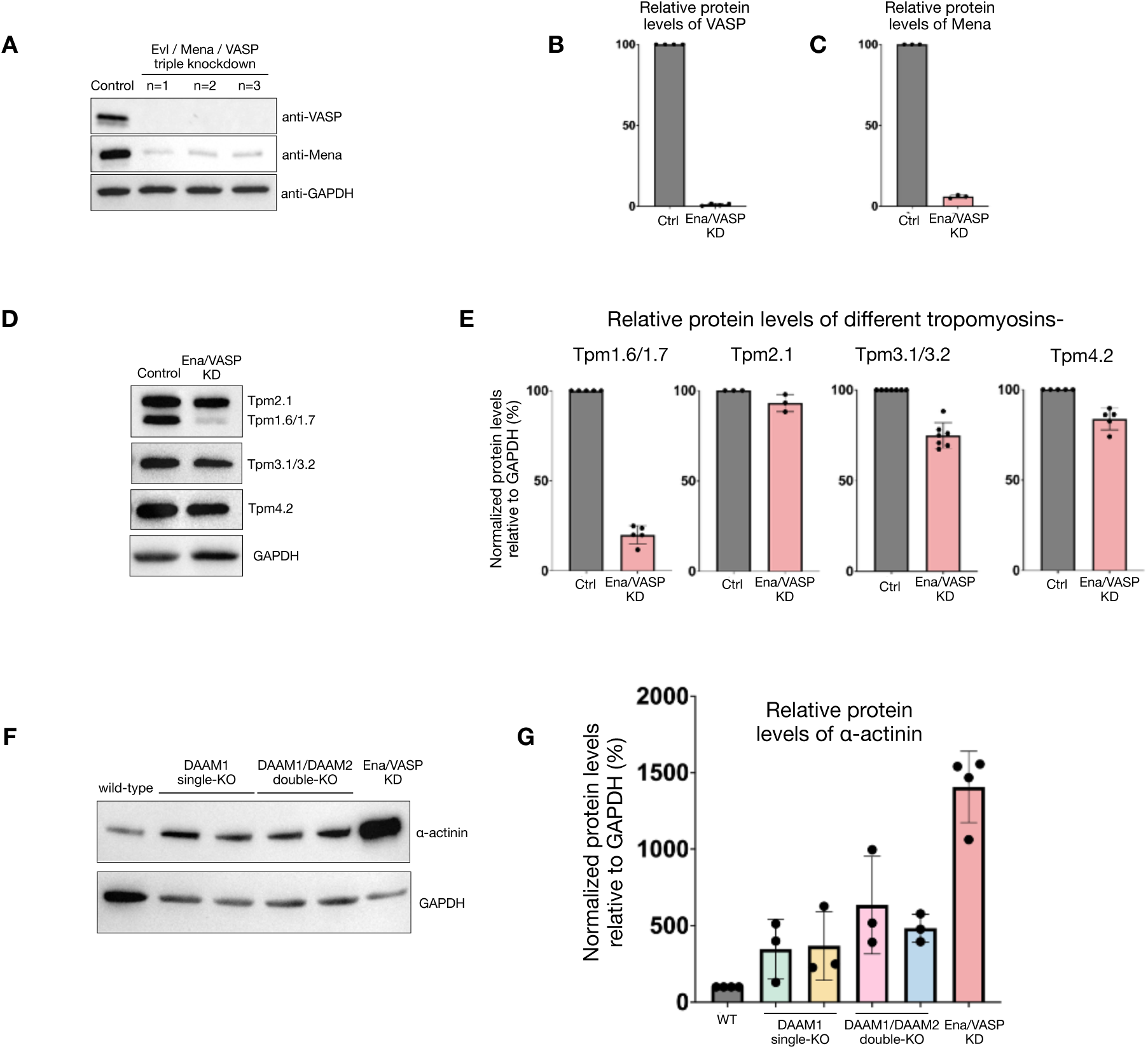
Relative protein levels of VASP, Mena, tropomyosins and α-actinin in Ena/VASP siRNA cells. **(A)** Western blot analysis and **(B-C)** quantification of the relative protein levels of VASP (panel B) and Mena (panel C) in control cells and cells treated with siRNA against Evl, Mena and VASP (Ena/VASP) for 72 hours from three different cell lysates. **(D)** Representative Western blots and **(E)** quantification of the relative protein levels of Tpm1.6/1.7 and Tpm2.1 (with TM311 antibody), Tpm3.1/3.2 (with γ9d antibody), and Tpm4.2 (with δ9d antibody) in the lysates of control and Ena/VASP- depleted cells. **(F)** Representative Western blots and **(G)** quantification of the relative protein levels of α-actinin in the lysates of wild-type, DAAM1 knockout clones #1 and #2, DAAM1/DAAM2 double knockout clones #1A and #1B, and Ena/VASP- depleted cells. Error bars represent mean ± SEM.

**Figure S5.**
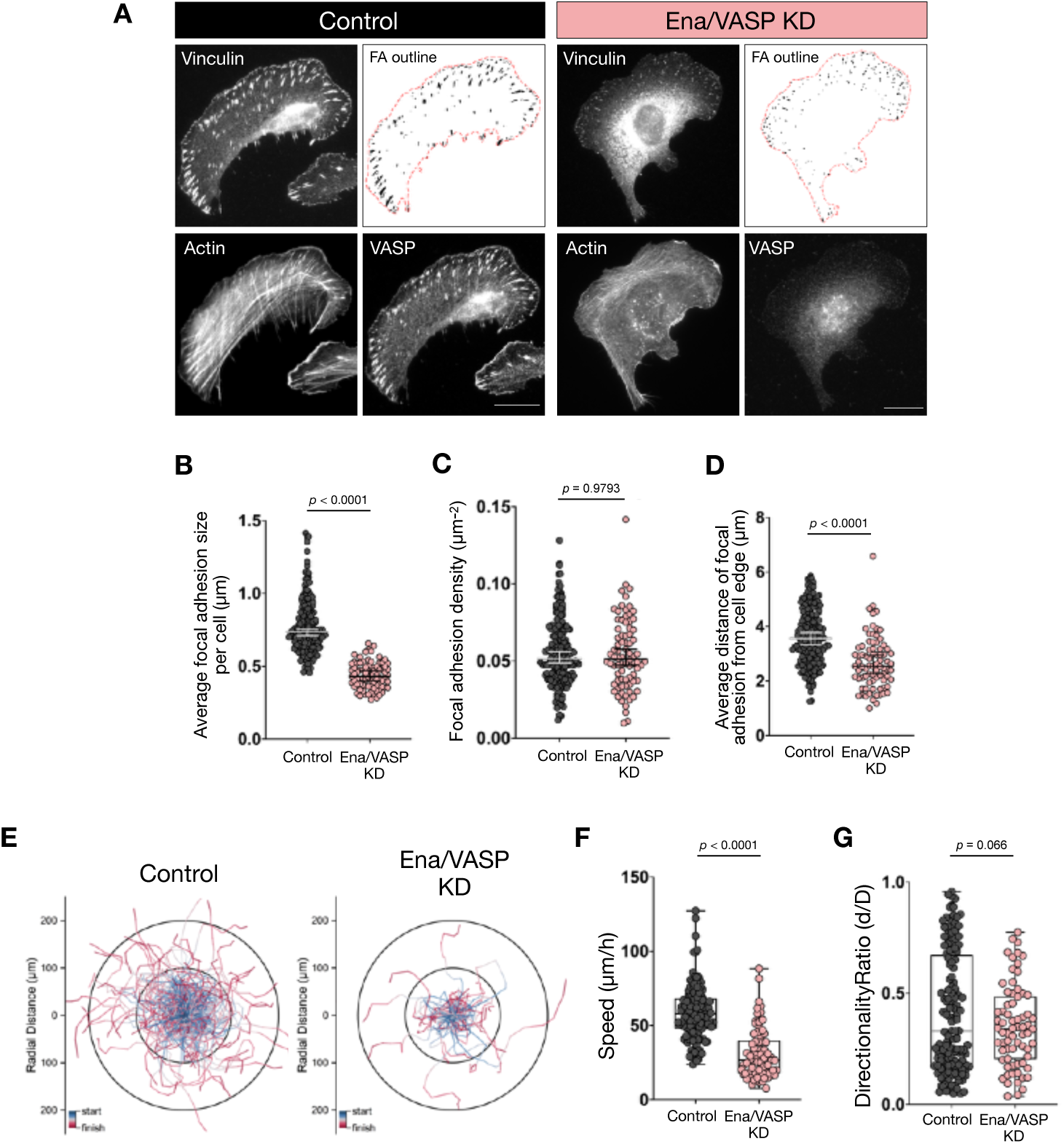
Phenotypes of Ena/VASP-depleted cells. Representative micrographs of control cells and cells treated with siRNAs against Evl, Mena and VASP (Ena/VASP) stained for focal adhesions (vinculin antibody), F-actin (phalloidin) and with anti-VASP antibody. Focal adhesion segmentation and cell outlines are also shown. **(B-D)** Quantification of the average focal adhesion size per cell (panel B), focal adhesion density (panel C), and average distance (µm) of focal adhesions from the cell edge (panel D) (*n* = 199 control cells, *n* = 80 Ena/VASP-depleted cells). **(E)** Random migration trajectories of control and Ena/VASP-depleted cells shown in polar coordinates with the start point synchronized at origin (0,0) and color-coded to indicate the track’s start (blue) and finish (red) after 5 h. **(F-G)** Quantification of cell migration speed (panel G) and directionality ratio (panel H). Each data point represents an individual cell. Error bars represent mean ± SEM.

**Figure S6.**
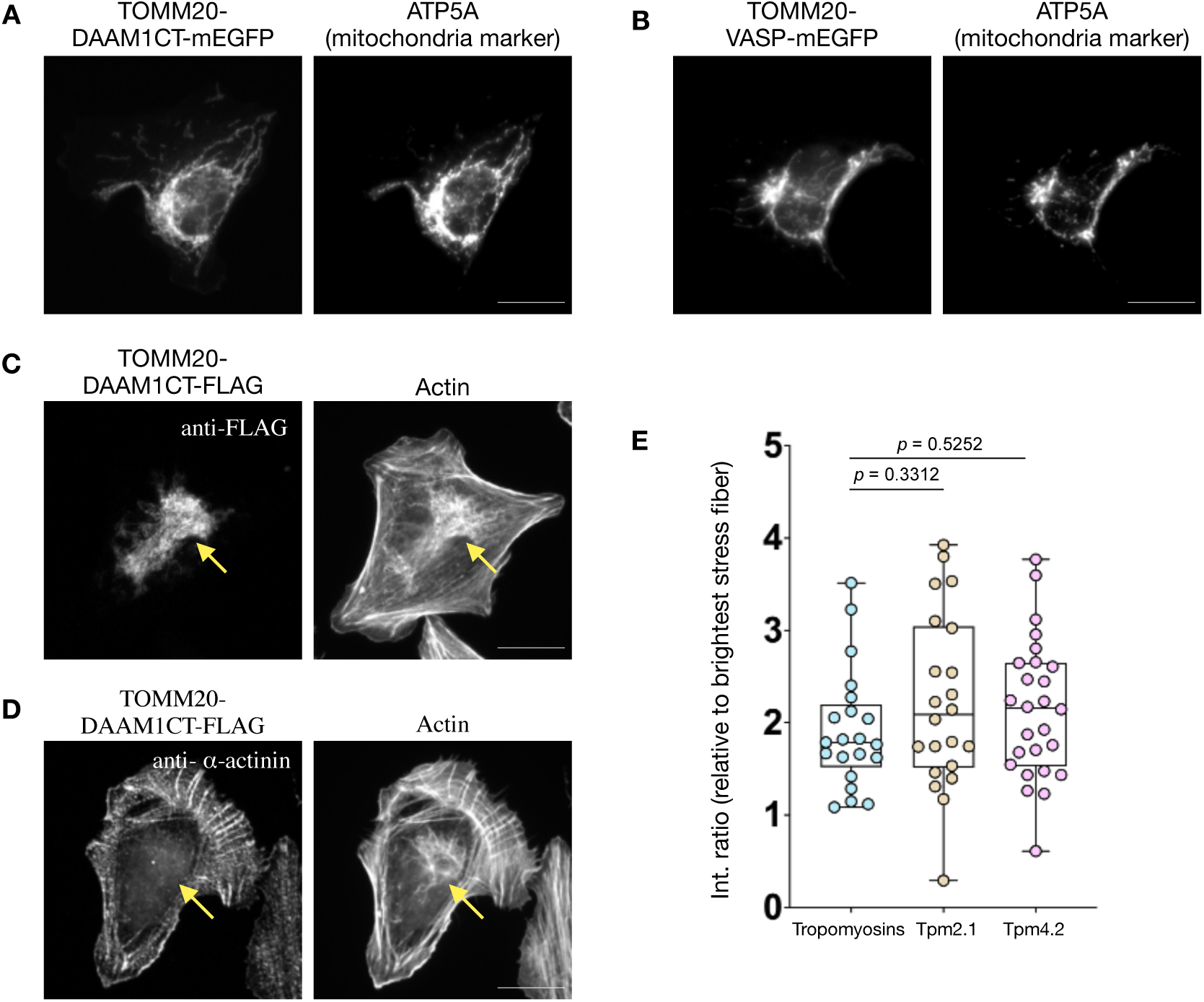
Validation of mitochondrial-targeted DAAM1 and VASP constructs. **(A-B)** Representative micrographs of cells expressing TOMM20-DAAM1CT-mEGFP (mito-DAAM1, panel A) and TOMM20- VASP-mEGFP (panel B) stained with mitochondria (ATP5A antibody). **(C-D)** Representative micrographs of cells expressing TOMM20-DAAM1CT-FLAG co-stained with phalloidin (for F-actin), and anti-FLAG (panel C) or anti-α-actinin (panel D) antibodies. **(E)** Quantification of the enrichment of endogenous tropomyosin signal at the mitochondria of cells expressing mito-DAAM1, in comparison to enrichment of mRuby-Tpm2.1 and mRuby-Tpm4.2 signal at the mitochondria. Error bars represent mean ± SEM. All scale bars, 20 µm.

**Figure S7.**
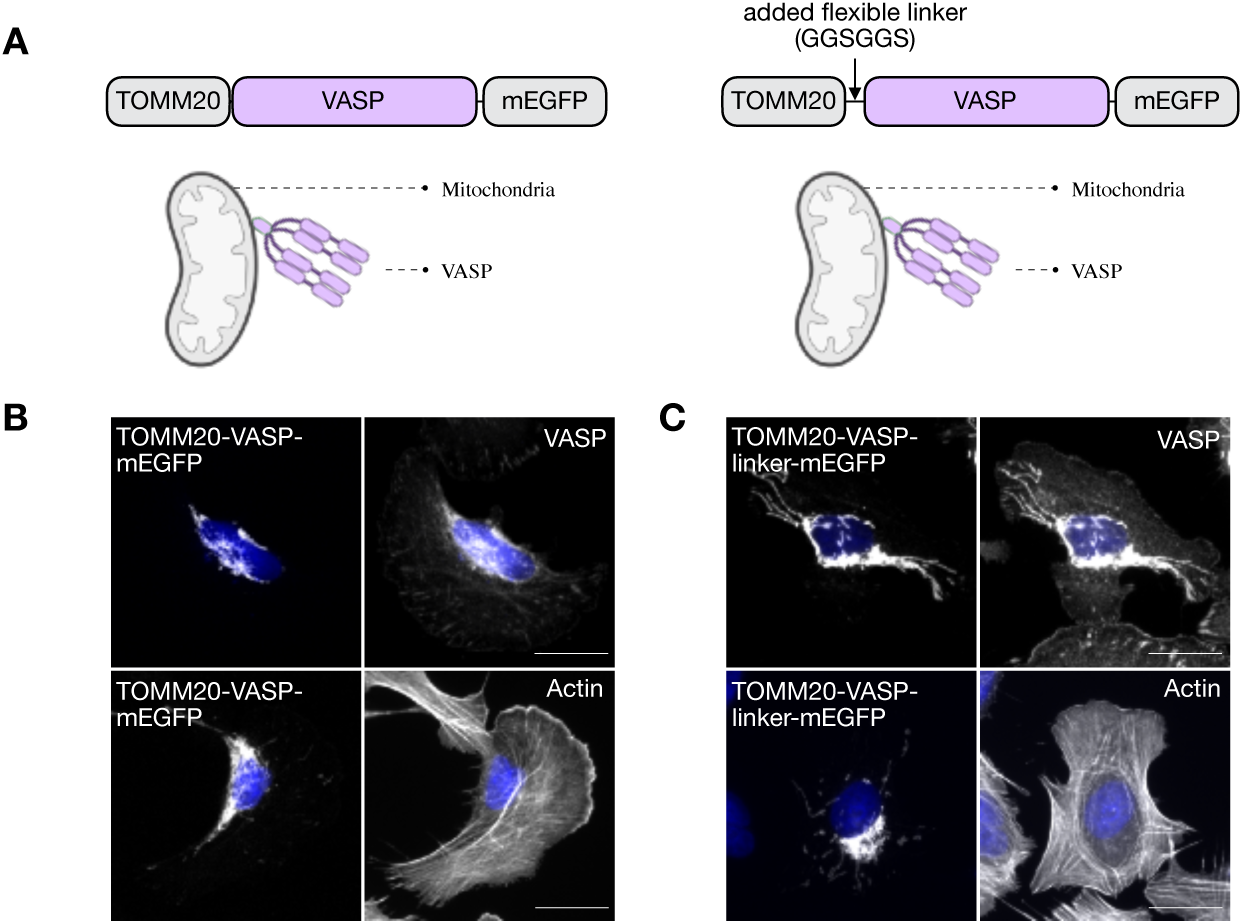
Mitochondrially-targeted full-length VASP constructs do not stimulate actin polymerization on the mitochondrial outer membrane. **(A)** Schematic representations of mitochondrially-targeted VASP constructs used in this study. **(B-C)** Representative micrographs of cells expressing TOMM20-VASP-mEGFP (panel B) and TOMM20-linker-VASP-mEGFP (panel C) co-stained with anti-VASP antibody and phalloidin (for F-actin). Scale bars, 20 µm.

**Figure S8.**
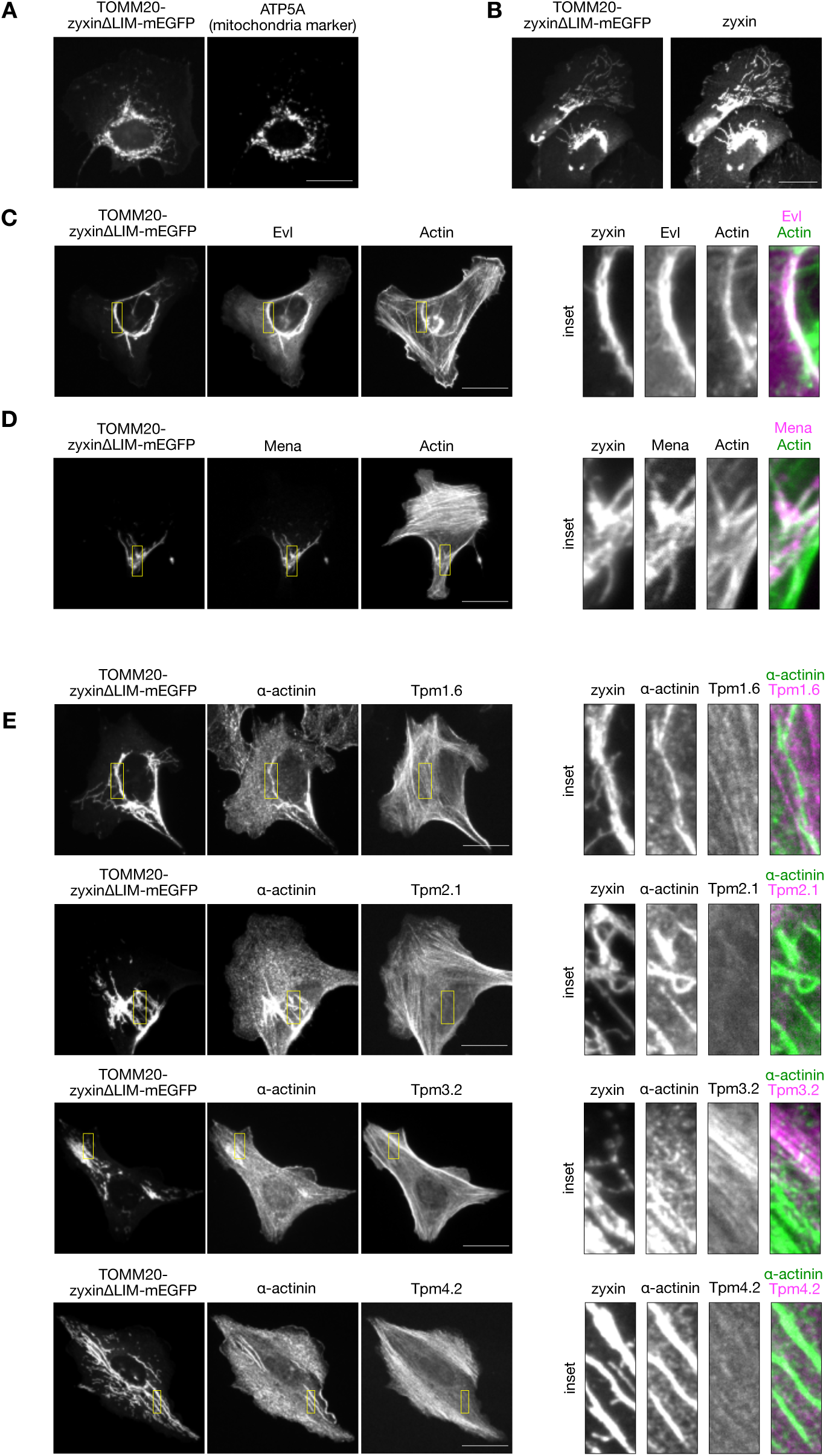
Mitochondrially-targeted zyxin recruits VASP, Evl and Mena, but does not induce the assembly of tropomyosin-decorated actin filaments on the mitochondrial outer membrane. **(A-B)** Representative micrographs of cells expressing TOMM20-zyxinΔLIM -mEGFP (mito-VASP) stained for mitochondria (ATP5A antibody, panel A) and with anti- zyxin antibody (panel B). **(C)** Representative micrographs of cells co-expressing mito-VASP and mScarlet3-Evl stained for F- actin (phalloidin). Yellow boxes in the whole cell images indicate the region shown in the insets on right. **(D)** Representative micrographs of cells expressing mito-VASP co-stained with anti-Mena antibody and phalloidin (for F-actin). Yellow boxes in the whole cell images indicate regions shown in the insets on right. **(E)** Representative micrographs of cells co-expressing mito-VASP and mRuby-Tpm1.6, -Tpm2.1, -Tpm3.2, or -Tpm4.2 and stained with antibody against α-actinin. Yellow boxes in the whole cell images indicate regions shown in the insets on right. All scale bars, 20 µm.

## References

1. Michelot, A., and Drubin, D.G. (2011). Building Distinct Actin Filament Networks in a Common Cytoplasm. Current Biology 21, R560–R569. 10.1016/j.cub.2011.06.019.

2. Pruyne, D.W., Schott, D.H., and Bretscher, A. (1998). Tropomyosin-containing Actin Cables Direct the Myo2p-dependent Polarized Delivery of Secretory Vesicles in Budding Yeast. J. Cell Biol. 143, 1931–1945. 10.1083/jcb.143.7.1931.

3. Kaksonen, M., Sun, Y., and Drubin, D.G. (2003). A Pathway for Association of Receptors, Adaptors, and Actin during Endocytic Internalization. Cell 115, 475–487. 10.1016/S0092-8674(03)00883-3.

4. Moseley, J.B., Maiti, S., and Goode, B.L. (2006). Formin Proteins: Purification and Measurement of Effects on Actin Assembly. In, pp. 215–234. 10.1016/S0076-6879(06)06016-2.

5. Mullins, R.D., Heuser, J.A., and Pollard, T.D. (1998). The interaction of Arp2/3 complex with actin: Nucleation, high affinity pointed end capping, and formation of branching networks of filaments. Proceedings of the National Academy of Sciences 95, 6181–6186. 10.1073/pnas.95.11.6181.

6. Pruyne, D., Evangelista, M., Yang, C., Bi, E., Zigmond, S., Bretscher, A., and Boone, C. (2002). Role of Formins in Actin Assembly: Nucleation and Barbed-End Association. Science. 297, 612– 615. 10.1126/science.1072309.

7. Sagot, I., Rodal, A.A., Moseley, J., Goode, B.L., and Pellman, D. (2002). An actin nucleation mechanism mediated by Bni1 and Profilin. Nat. Cell Biol. 4, 626–631. 10.1038/ncb834.

8. Palmer, N.J., Barrie, K.R., and Dominguez, R. (2024). Mechanisms of actin filament severing and elongation by formins. Nature 632, 437–442. 10.1038/s41586-024-07637-0.

9. Oosterheert, W., Boiero Sanders, M., Funk, J., Prumbaum, D., Raunser, S., and Bieling, P. (2024). Molecular mechanism of actin filament elongation by formins. Science 384, eadn9560. 10.1126/science.adn9560.

10. Skau, C.T., and Kovar, D.R. (2010). Fimbrin and Tropomyosin Competition Regulates Endocytosis and Cytokinesis Kinetics in Fission Yeast. Current Biology 20, 1415–1422. 10.1016/j.cub.2010.06.020.

11. Kadzik, R.S., Homa, K.E., and Kovar, D.R. (2020). F-Actin Cytoskeleton Network Self-Organization Through Competition and Cooperation. Annu. Rev. Cell Dev. Biol. 36, 35–60. 10.1146/annurev-cellbio-032320-094706.

12. Homa, K.E., Hocky, G.M., Suarez, C., and Kovar, D.R. (2024). Arp2/3 complex- and formin-mediated actin cytoskeleton networks facilitate actin binding protein sorting in fission yeast. Eur. J. Cell Biol. 103, 151404. 10.1016/j.ejcb.2024.151404.

13. Burke, T.A., Christensen, J.R., Barone, E., Suarez, C., Sirotkin, V., and Kovar, D.R. (2014). Homeostatic Actin Cytoskeleton Networks Are Regulated by Assembly Factor Competition for Monomers. Current Biology 24, 579–585. 10.1016/j.cub.2014.01.072.

14. Rotty, J.D., Wu, C., Haynes, E.M., Suarez, C., Winkelman, J.D., Johnson, H.E., Haugh, J.M., Kovar, D.R., and Bear, J.E. (2015). Profilin-1 Serves as a Gatekeeper for Actin Assembly by Arp2/3-Dependent and -Independent Pathways. Dev. Cell 32, 54–67. 10.1016/j.devcel.2014.10.026.

15. Suarez, C., Carroll, R.T., Burke, T.A., Christensen, J.R., Bestul, A.J., Sees, J.A., James, M.L., Sirotkin, V., and Kovar, D.R. (2015). Profilin Regulates F-Actin Network Homeostasis by Favoring Formin over Arp2/3 Complex. Dev. Cell 32, 43–53. 10.1016/j.devcel.2014.10.027.

16. Vitriol, E.A., McMillen, L.M., Kapustina, M., Gomez, S.M., Vavylonis, D., and Zheng, J.Q. (2015). Two Functionally Distinct Sources of Actin Monomers Supply the Leading Edge of Lamellipodia. Cell Rep. 11, 433–445. 10.1016/j.celrep.2015.03.033.

17. Chua, X. Le, Tong, C.S., Su, M., Xŭ, X.J., Xiao, S., Wu, X., and Wu, M. (2024). Competition and synergy of Arp2/3 and formins in nucleating actin waves. Cell Rep. 43, 114423. 10.1016/j.celrep.2024.114423.

18. Blanchoin, L., Boujemaa-Paterski, R., Sykes, C., and Plastino, J. (2014). Actin Dynamics, Architecture, and Mechanics in Cell Motility. Physiol. Rev. 94, 235–263. 10.1152/physrev.00018.2013.

19. Lappalainen, P., Kotila, T., Jégou, A., and Romet-Lemonne, G. (2022). Biochemical and mechanical regulation of actin dynamics. Nat. Rev. Mol. Cell Biol. 23, 836–852. 10.1038/s41580-022-00508-4.

20. Hardeman, E.C., Bryce, N.S., and Gunning, P.W. (2020). Impact of the actin cytoskeleton on cell development and function mediated via tropomyosin isoforms. Semin. Cell Dev. Biol. 102, 122– 131. 10.1016/j.semcdb.2019.10.004.

21. Gateva, G., Kremneva, E., Reindl, T., Kotila, T., Kogan, K., Gressin, L., Gunning, P.W., Manstein, D.J., Michelot, A., and Lappalainen, P. (2017). Tropomyosin Isoforms Specify Functionally Distinct Actin Filament Populations In Vitro. Current Biology 27, 705–713. 10.1016/j.cub.2017.01.018.

22. Higgs, H.N. (2005). Formin proteins: a domain-based approach. Trends Biochem. Sci. 30, 342– 353. 10.1016/j.tibs.2005.04.014.

23. Faix, J., and Rottner, K. (2022). Ena/VASP proteins in cell edge protrusion, migration and adhesion. J. Cell Sci. 135, jcs259226. 10.1242/jcs.259226.

24. Kumari, R., Ven, K., Chastney, M., Kokate, S.B., Peränen, J., Aaron, J., Kogan, K., Almeida-Souza, L., Kremneva, E., Poincloux, R., et al. (2024). Focal adhesions contain three specialized actin nanoscale layers. Nat. Commun. 15, 2547. 10.1038/s41467-024-46868-7.

25. Tojkander, S., Gateva, G., Schevzov, G., Hotulainen, P., Naumanen, P., Martin, C., Gunning, P.W., and Lappalainen, P. (2011). A Molecular Pathway for Myosin II Recruitment to Stress Fibers. Current Biology 21, 539–550. 10.1016/j.cub.2011.03.007.

26. Ang, S.F., Zhao, Z.S., Lim, L., and Manser, E. (2010). DAAM1 is a formin required for centrosome re-orientation during cell migration. PLoS One 5(9):e13064. 10.1371/journal.pone.0013064.

27. Yan, T., Zhang, A., Shi, F., Chang, F., Mei, J., Liu, Y., and Zhu, Y. (2018). Integrin ávâ3– associated DAAM1 is essential for collagen-induced invadopodia extension and cell haptotaxis in breast cancer cells. Journal of Biological Chemistry 293, 10172–10185. 10.1074/jbc.RA117.000327.

28. Luo, W., Lieu, Z.Z., Manser, E., Bershadsky, A.D., and Sheetz, M.P. (2016). Formin DAAM1 Organizes Actin Filaments in the Cytoplasmic Nodal Actin Network. PLoS One 11, e0163915. 10.1371/journal.pone.0163915.

29. Sherrard, K.M., Cetera, M., and Horne-Badovinac, S. (2021). DAAM mediates the assembly of long-lived, treadmilling stress fibers in collectively migrating epithelial cells in Drosophila. Elife 10:e72881. 10.7554/eLife.72881.

30. Meiring, J.C.M., Bryce, N.S., Wang, Y., Taft, M.H., Manstein, D.J., Liu Lau, S., Stear, J., Hardeman, E.C., and Gunning, P.W. (2018). Co-polymers of Actin and Tropomyosin Account for a Major Fraction of the Human Actin Cytoskeleton. Current Biology 28, 2331–2337.e5. 10.1016/j.cub.2018.05.053.

31. Kumari, R., Jiu, Y., Carman, P.J., Tojkander, S., Kogan, K., Varjosalo, M., Gunning, P.W., Dominguez, R., and Lappalainen, P. (2020). Tropomodulins Control the Balance between Protrusive and Contractile Structures by Stabilizing Actin-Tropomyosin Filaments. Current Biology 30, 767–778.e5. 10.1016/j.cub.2019.12.049.

32. Bear, J.E., Loureiro, J.J., Libova, I., Fässler, R., Wehland, J., and Gertler, F.B. (2000). Negative Regulation of Fibroblast Motility by Ena/VASP Proteins. Cell 101, 717–728. 10.1016/S0092-8674(00)80884-3.

33. Chen, X.J., Squarr, A.J., Stephan, R., Chen, B., Higgins, T.E., Barry, D.J., Martin, M.C., Rosen, M.K., Bogdan, S., and Way, M. (2014). Ena/VASP Proteins Cooperate with the WAVE Complex to Regulate the Actin Cytoskeleton. Dev. Cell 30, 569–584. 10.1016/j.devcel.2014.08.001.

34. Gateva, G., Tojkander, S., Koho, S., Carpen, O., and Lappalainen, P. (2014). Palladin promotes assembly of non-contractile dorsal stress fibers through VASP recruitment. J. Cell Sci. 127(9): 1887–1898. 10.1242/jcs.135780.

35. Damiano-Guercio, J., Kurzawa, L., Mueller, J., Dimchev, G., Schaks, M., Nemethova, M., Pokrant, T., Brühmann, S., Linkner, J., Blanchoin, L., et al. (2020). Loss of Ena/VASP interferes with lamellipodium architecture, motility and integrin-dependent adhesion. Elife 9:e55351. 10.7554/eLife.55351.

36. Hotulainen, P., and Lappalainen, P. (2006). Stress fibers are generated by two distinct actin assembly mechanisms in motile cells. J. Cell Biol. 173, 383–394. 10.1083/jcb.200511093.

37. Kanchanawong, P., Shtengel, G., Pasapera, A.M., Ramko, E.B., Davidson, M.W., Hess, H.F., and Waterman, C.M. (2010). Nanoscale architecture of integrin-based cell adhesions. Nature 468, 580–584. 10.1038/nature09621.

38. Vartiainen, M.K., Guettler, S., Larijani, B., and Treisman, R. (2007). Nuclear Actin Regulates Dynamic Subcellular Localization and Activity of the SRF Cofactor MAL. Science. 316, 1749– 1752. 10.1126/science.1141084.

39. Sun, Q., Chen, G., Streb, J.W., Long, X., Yang, Y., Stoeckert, C.J., and Miano, J.M. (2006). Defining the mammalian CArGome. Genome Res. 16, 197–207. 10.1101/gr.4108706.

40. Gau, D., and Roy, P. (2018). SRF’ing and SAP’ing – the role of MRTF proteins in cell migration. J. Cell Sci. 131, jcs218222. 10.1242/jcs.218222.

41. Grosse, R., Copeland, J.W., Newsome, T.P., Way, M., and Treisman, R. (2003). A role for VASP in RhoA-Diaphanous signalling to actin dynamics and SRF activity. EMBO J. 22, 3050–3061. 10.1093/emboj/cdg287.

42. Hoffman, L.M., Jensen, C.C., Kloeker, S., Wang, C.-L.A., Yoshigi, M., and Beckerle, M.C. (2006). Genetic ablation of zyxin causes Mena/VASP mislocalization, increased motility, and deficits in actin remodeling. J. Cell Biol. 172, 771–782. 10.1083/jcb.200512115.

43. Winkelman, J.D., Anderson, C.A., Suarez, C., Kovar, D.R., and Gardel, M.L. (2020). Evolutionarily diverse LIM domain-containing proteins bind stressed actin filaments through a conserved mechanism. Proceedings of the National Academy of Sciences 117, 25532–25542. 10.1073/pnas.2004656117.

44. Schevzov, G., Whittaker, S.P., Fath, T., Lin, J.J.-C., and Gunning, P.W. (2011). Tropomyosin isoforms and reagents. Bioarchitecture 1, 135–164. 10.4161/bioa.1.4.17897.

45. Case, L.B., and Waterman, C.M. (2015). Integration of actin dynamics and cell adhesion by a three-dimensional, mechanosensitive molecular clutch. Nat. Cell Biol. 17, 955–963. 10.1038/ncb3191.

46. Stubb, A., Guzmán, C., Närvä, E., Aaron, J., Chew, T.-L., Saari, M., Miihkinen, M., Jacquemet, G., and Ivaska, J. (2019). Superresolution architecture of cornerstone focal adhesions in human pluripotent stem cells. Nat. Commun. 10, 4756. 10.1038/s41467-019-12611-w.

47. Lehtimäki, J.I., Rajakylä, E.K., Tojkander, S., and Lappalainen, P. (2021). Generation of stress fibers through myosin-driven reorganization of the actin cortex. Elife 10:e60710. 10.7554/eLife.60710.

48. Brühmann, S., Ushakov, D.S., Winterhoff, M., Dickinson, R.B., Curth, U., and Faix, J. (2017). Distinct VASP tetramers synergize in the processive elongation of individual actin filaments from clustered arrays. Proceedings of the National Academy of Sciences 114 (29) E815–E5824. 10.1073/pnas.1703145114.

49. Winkelman, J.D., Bilancia, C.G., Peifer, M., and Kovar, D.R. (2014). Ena/VASP Enabled is a highly processive actin polymerase tailored to self-assemble parallel-bundled F-actin networks with Fascin. Proceedings of the National Academy of Sciences 111, 4121–4126. 10.1073/pnas.1322093111.

50. Winkelman, J.D., Suarez, C., Hocky, G.M., Harker, A.J., Morganthaler, A.N., Christensen, J.R., Voth, G.A., Bartles, J.R., and Kovar, D.R. (2016). Fascin- and α-Actinin-Bundled Networks Contain Intrinsic Structural Features that Drive Protein Sorting. Current Biology 26, 2697–2706. 10.1016/j.cub.2016.07.080.

51. Kovar, D.R. (2006). Molecular details of formin-mediated actin assembly. Curr. Opin. Cell Biol. 18, 11–17. 10.1016/j.ceb.2005.12.011.

52. Fowler, V.M., and Dominguez, R. (2017). Tropomodulins and Leiomodins: Actin Pointed End Caps and Nucleators in Muscles. Biophys. J. 112, 1742–1760. 10.1016/j.bpj.2017.03.034.

53. Carman, P.J., Barrie, K.R., Rebowski, G., and Dominguez, R. (2023). Structures of the free and capped ends of the actin filament. Science 380, 1287–1292. 10.1126/science.adg6812.

54. Kovar, D.R., Harris, E.S., Mahaffy, R., Higgs, H.N., and Pollard, T.D. (2006). Control of the Assembly of ATP- and ADP-Actin by Formins and Profilin. Cell 124, 423–435. 10.1016/j.cell.2005.11.038.

55. Vavylonis, D., Kovar, D.R., O’Shaughnessy, B., and Pollard, T.D. (2006). Model of Formin-Associated Actin Filament Elongation. Mol. Cell 21, 455–466. 10.1016/j.molcel.2006.01.016.

56. Funk, J., Merino, F., Venkova, L., Heydenreich, L., Kierfeld, J., Vargas, P., Raunser, S., Piel, M., and Bieling, P. (2019). Profilin and formin constitute a pacemaker system for robust actin filament growth. Elife 8:e50963. 10.7554/eLife.50963.

57. 57. Guardia, C.M., De Pace, R., Sen, A., Saric, A., Jarnik, M., Kolin, D.A., Kunwar, A., and Bonifacino, J.S. (2019). Reversible association with motor proteins (RAMP): A streptavidin-based method to manipulate organelle positioning. PLoS Biol. 17, e3000279. 10.1371/journal.pbio.3000279.

58. Spudich, J.A., and Watt, S. (1971). The regulation of rabbit skeletal muscle contraction. I. Biochemical studies of the interaction of the tropomyosin-troponin complex with actin and the proteolytic fragments of myosin. J. Biol. Chem. 246, 4866–4871.

59. Wioland, H., Guichard, B., Senju, Y., Myram, S., Lappalainen, P., Jégou, A., and Romet-Lemonne, G. (2017). ADF/Cofilin Accelerates Actin Dynamics by Severing Filaments and Promoting Their Depolymerization at Both Ends. Current Biology 27, 1956–1967.e7. 10.1016/j.cub.2017.05.048.

60. Bagès, C., Chabanon, M., Kools, W., Dos Santos, T., Pagès, R., Sirkia, M.E., Leduc, C., Houdusse, A., Jégou, A., Romet-Lemonne, G., et al. (2025). Probing protein–protein interactions with drag flow: a case study of F-actin and tropomyosin. The European Physical Journal E 48, 49. 10.1140/epje/s10189-025-00509-z.

61. Kokate, S.B., Oshin, A.T., Chua, X. Le, Chastney, M., Biswas, P., Kogan, K., Tomberg, T., Ivaska, J., and Lappalainen, P. (2025). Calponin isoforms define the cell-type-specific organization and dynamics of actomyosin bundles. Current Biology 35, 6024–6037.e8. 10.1016/j.cub.2025.10.081.

62. Ran, F.A., Hsu, P.D., Wright, J., Agarwala, V., Scott, D.A., and Zhang, F. (2013). Genome engineering using the CRISPR-Cas9 system. Nat. Protoc. 8, 2281–2308. 10.1038/nprot.2013.143.

63. Seetharaman, S., Vianay, B., Roca, V., Farrugia, A.J., De Pascalis, C., Boëda, B., Dingli, F., Loew, D., Vassilopoulos, S., Bershadsky, A., et al. (2022). Microtubules tune mechanosensitive cell responses. Nat. Mater. 21, 366–377. 10.1038/s41563-021-01108-x.

